# Exploring the conformational landscape and stability of Aurora A using ion-mobility mass spectrometry and molecular modelling

**DOI:** 10.1101/2021.08.30.458190

**Authors:** Lauren J. Tomlinson, Matthew Batchelor, Joscelyn Sarsby, Dominic P. Byrne, Philip Brownridge, Richard Bayliss, Patrick A. Eyers, Claire E. Eyers

**Author notes:** Correspondence: Claire E. Eyers.

## Abstract

Protein kinase inhibitors are proving highly effective in helping treat a number of non-communicable diseases driven by aberrant kinase signaling. They are also extremely valuable as chemical tools to help delineate cellular roles of kinase signaling complexes. The binding of small molecule inhibitors induces conformational effects on kinase dynamics; evaluating the effect of such interactions can assist in developing specific inhibitors and is deemed imperative to understand both inhibition and resistance mechanisms. Using gas-phase ion mobility-mass spectrometry (IM-MS) we characterized changes in the conformational landscape and stability of the protein kinase Aurora A (Aur A) driven by binding of the physiological activator TPX2 or small molecule inhibition. Aided by molecular modeling, we establish three major conformations: one highly-populated compact conformer similar to that observed in most crystal structures, a second highly-populated conformer possessing a more open structure that is infrequently found in crystal structures, and an additional low-abundance conformer not currently represented in the protein databank. Comparison of active (phosphorylated) and inactive (non-phosphorylated) forms of Aur A revealed that the active enzyme has different conformer weightings and is less stable than the inactive enzyme. Notably, inhibitor binding shifts conformer balance towards the more compact configurations adopted by the unbound enzyme, with both IM-MS and modelling revealing inhibitor-mediated stabilisation of active Aur A. These data highlight the power of IM-MS in combination with molecular dynamics simulations to probe and compare protein kinase structural dynamics that arise due to differences in activity and as a result of compound binding.

## Introduction

Protein kinase-mediated phosphorylation permits dynamic regulation of protein function and is an essential mechanism for modulating a host of fundamental biological processes. Inhibition of these enzymes by small molecules can serve both as a research tool to help understand cell signalling mechanisms, and also to help treat diseases such as cancer, inflammatory disorders and diabetes (1), where protein phosphorylation is often dysregulated.

Protein kinases consist of an N-terminal and a C-terminal lobe, connected via a flexible hinge region, which forms the conserved ATP-binding site. The activation loop in protein kinases is 20–30 residues long with a conserved DFG motif at the beginning, typically extending out to an invariant APE motif. Activation loops of active kinases are mobile, and help form a cleft that enables substrates to bind. Substrates are then positioned adjacent to the HRD motif, where the catalytic Asp residue acts as the catalytic base to permit productive phosphoryation (2).

The ATP-binding sites of members of the kinase superfamily share a high degree of sequence conservation, which has made the development of highly selective ATP-competitive inhibitor compounds extremely challenging, and usually requires the exploitation of unique, often subtle, structural deviations in individual enzymes. Understanding the selectivity and specificity of these small molecule inhibitors towards target enzymes is critical for correct interpretation of data arising from their use, due to the likelihood of ‘off-target’ effects driven through similar kinase conformations that exist across members of the evolutionary-related kinome (3).

Protein kinase inhibitors can be broadly classified based on their ability to bind to different regions within the enzyme superfamily, or to a specific conformational state. In catalytically active kinases, the activation loop usually exists in a ‘DFG-in’ conformation, orientating the conserved DFG motif to support metal binding as the DFG-Phe contacts the C-helix of the N-terminal lobe. In contrast, a kinase occupying a ‘DFG-out’ conformation is less able to bind ATP, since the Phe partially occludes the metal:nucleotide binding site and exposes a C-helix pocket, rendering it catalytically inactive (4). The majority of kinase small molecule inhibitors function by disrupting the ability of kinases to bind to- and/or hydrolyse ATP and therefore block phosphate transfer to protein substrates, either by competing directly with ATP binding, or by locking the enzyme in an ‘inactive’ conformation.

Critically, much of our current understanding of kinase inhibitor binding modes comes from X-ray crystallography of kinase:inhibitor complexes. While ‘type I’ inhibitors, such as staurosporine and dasatanib, competitively bind to the ATP-binding site of kinases in the active ‘DFG-in’ conformation, ‘type II’ inhibitors like imatinib are ‘mixed mode’, contacting both the ATP binding site and an adjacent hydrophobic groove that is only accessible in the ‘DFG-out’ conformation, which serves to lock the target kinase into an inactive state (5). The ability of small molecules to discriminate between, and selectively bind to, kinases in various active and inactive structural orientations has been used as a defining tool to group classes of inhibitor compounds (6). However, in addition to significant diversity in the DFG-in and (in particular) DFG-out structures of multiple kinases, a variety of kinases exist in which the Phe residue of the DFG motif can adopt an ‘intermediary’ orientation between the typical DFG-in and DFG-out conformation, termed DFG-up or DFG-inter (2).

Characterizing the effects of small molecule inhibitors on the structure and catalytic activity of protein kinase targets, and the influence of post-translational modifications (PTMs; typically activating phosphorylation) on these interactions provides fundamental mechanistic knowledge for drug discovery and helps iterative drug design. However, limitations often arise with crystallographic structural studies, with some protein being intransigent to crystallisation (7). Moreover, the analysis of solid-state crystals hampers the ability to define and understand conformational flexibility and protein dynamics in both the absence and presence of bound small molecules.

Consequently, analysis of protein crystal structures may only permit limited exploration of the conformational space adopted by kinases in solution. NMR, while useful for examining conformational dynamics of purified proteins, requires much more significant (multi-milligram) amounts of pure material, and obtaining a full atomic map for larger proteins or complexes greater than ∼50 kDa remains a challenge (8). Native mass spectrometry (MS), in which liquid-phase samples are subjected to electrospray ionisation (ESI) under non-denaturing conditions to more closely mimic their physiological environment, is increasingly being used to investigate the topology of intact protein complexes (9). Under carefully controlled conditions (pH, ionic strength, applied voltage, gas-pressure), the folded native state of the analyte protein (and ligand) complexes can be maintained (10). Native MS is primarily used to define the molecular mass of protein complexes (and component stoichiometry), compare the relative dissociation contant (*K*_D_) of ligand binding (11-14), and in a broad sense, the degree of protein ‘disorder’ (15). When used in combination with ion mobility (IM) spectrometry, native MS (IM-MS) can reveal structural changes that arise due to ligand binding or protein modification as well as interrogating protein conformational dynamics, stability and unfolding transitions (16). When appropriately calibrated, native IM-MS can also be used to determine the rotationally averaged collision cross-section (CCS) of proteins and their complexes, empirical information that can be compared both with other structural measurements, and theoretical calculations (13) to understand the effects of PTMs or small molecule binding on protein structure and dynamics.

The Ser/Thr protein kinase Aurora A (Aur A) is classically associated with mitotic entry and plays critical roles in centrosome maturation and separation (17). In early G phase of the cell cycle, Aur A is recruited to the centrosomes where it facilitates spindle assembly (18, 19). Later, Aur A microtubule association and activation requires binding of the Eg5-associated microtubule factor TPX2 (20-22). The N-terminus of TPX2 binds to the Aur A catalytic domain, inducing conformational changes in the kinase which both enhances its autophosphorylation at Thr288 within the activation loop, and shields the activating T-loop phosphorylation site from phosphatases such as PP6 (23-25). Overexpression of Aur A results in mitotic abnormalities and the development of tetraploid cells (26). While elevated levels of Aur A are broadly associated with a range of cancers, including breast, colorectal, ovarian and pancreatic (27), depletion of Aur A activity leads to abnormalities in mitotic spindle assembly, which results in a spindle checkpoint-dependent mitotic arrest (28-31). More recently, catalytically-independent roles for Aur A in different phases of the cell cycle have also been described (30, 32-35).

A huge number of Aur A inhibitors have been developed and reported in the last two decades (36), many of which target all three members of the Aurora kinase family, most notably the well-studied pan Aur inhibitor, VX-680/tozasertib (31, 37). Inhibitors that show a preference between human Aurora kinases have also been developed by targeting specific amino acid differences in the ATP site that occur between Aur A and Aur B/C (38-40). For example, Alisterib (MLN8237) is a selective Aur A inhibitor that alongside the earlier tool compound MLN8054 (41, 42) has been characterised using a variety of *in vitro* and *in vivo* pre-clinical models (43). Importantly, MLN8237 has been phenotypically target-validated in cells with drug-resistant Aur A alleles (34, 42). It has also been shown to inhibit proliferation in a number of human tumour cell lines, such as ovarian, prostate, lung and lymphoma cells, and has proven effective in paediatric-type cancer models, including neuroblastoma, acute lymphoblastic leukaemia and Ewing sarcoma cell lines (44). MLN8237 has been assessed in Phase I and II clinical trials for haematological malignancies, patients with advanced solid tumours and children with refractory/recurrent solid tumours (45, 46).

To better understand the effects of small molecule binding on Aur A, Levinson and colleagues recently reported a study to evaluate the conformational effects of a panel of clinically relevent Aur kinase inhibitors across different activation states of Aur A using time-resolved Förster resonance energy transfer (TR-FRET). Using this approach, they were able to track dynamic structural movements of the kinase activation loop, distinguishing between inhibitors that induce DFG-in states from compounds that promote other conformations (DFG-out/DFG-up/DFG-inter). The TR-FRET data was consistent with equilibrium shifts towards three distinct conformational groups, including DFG-in state, DFG-out state and ‘unique’ structural states (47).

In this study, we employ IM-MS to explore the effects of inhibitor binding on the conformational landscape, dynamics and stability of two variants of the Aur A kinase domain; a catalytically active hyperphosphorylated protein, and an inactive non-phosphorylated version created by a point mutation within the DFG motif (D274N). These studies reveal marked differences in the conformational landscape adopted by active and inactive Aur A and also when complexed with inhibitors, with the active form presenting a shift in the conformer balance towards a less conformationally flexible configuration. Crucially, our CIU data also suggest that chemical inhibitors induce stabilisation of both hyper- and non-phosphorylated Aur A, whilst revealing intermediate unfolding transition differences that correlate with previously reported DFG in/out/up classifications with distinct compounds. Together, our biophysical data demonstrate the applicability of IM-MS for distinguishing modes of inhibitor binding to kinases that could be extendable to other members of the highly druggable superfamily.

## Materials and Methods

### Protein purification

6His-N-terminally tagged human Aur A (122-403) wild-type (WT) or D274N were individually expressed from a pET30-TEV vector in BL21 (DE3) pLysS *Escherichia coli* (Novagen), with protein expression being induced with 0.4 mM IPTG for 18 h at 18 °C. *E. coli* pellets were lysed in 100 mL of ice cold lysis buffer (50 mM Tris-HCl pH 7.4, 10% glycerol, 300 mM NaCl, 10 mM imidazole, 1 mM DTT, 100 mM EDTA, 100 mM EGTA, protease inhibitor tablet (Roche)). The lysed cells were then sonicated on ice using a 3 mm microprobe attached to a MSE Soniprep 150 plus motor unit at an amplitude of 16 microns in 30 second intervals. Samples were centrifuged for 1 h at 8 °C (43,000 *x g*) to pellet the cellular debris and then filtered through a 0.22 µm filter. His-tagged Aur A was separated from clarified bacterial cell lysate using a Nickel HisTrap HP column, pre-equilibrated in wash buffer (50 mM Tris-HCI pH 7.0, 10% glycerol, 300 mM NaCl, 20 mM imidazole, 1 mM MgCl_2_). After loading the cell lysate, the column was washed with 10 mL of wash buffer, followed by 10 mL of elution buffer (50 mM Tris-HCI pH 7.0, 10% glycerol, 300 mM NaCl, 500 mM imidazole, 1 mM MgCl_2_, 1 mM DTT) and the His-tag cleaved by addition of 25 µg of TEV protease and incubation for 18 h at 4 °C. Subsequently, Aur A was further purified using a Superdex 200 16 600 column (GE Healthcare) attached to an AKTA FPLC system and a Frac-920 (GE Healthcare), which was equilibrated in filtered and degassed gel filtration buffer (20 mM Tris pH 7.0, 10% glycerol, 200 mM NaCl, 40 mM imidazole, 5 mM MgCl_2_, 1 mM DTT). Aur A-containing fractions were pooled and passed through a HisTrap column to remove residual non-TEV cleaved material. Samples were stored in small aliquots at -80 °C prior to further analysis.

### Native Ion Mobility-Mass Spectrometry

Immediately prior to native MS analysis, purified Aur A proteins were buffer-exchanged into 150 mM NH_4_OAc using an Amicon spin filter (10 kDa cut-off). Spin columns were pre-washed with 500 µL of 150 mM NH_4_OAc prior to the addition of protein and spun 3x 10 min at 13,000 RPM. Following the final spin, the filter was inverted into a new collection tube and spun for 2 min at 3,000 RPM to collect the protein. Protein concentration was calculated using a NanoDrop spectrophotometer at a wavelength of 280 nm and adjusted to 5 µM for MS analysis. To evaluate the effect of small molecule binding, Aur A proteins were incubated with 4% DMSO (vehicle control), or a 10x molar excess of inhibitor or TPX2 activating 43mer peptide (H_2_N-MSQVKSSYSYDAPSDFINFSSLDDEGDTQNIDSWFEEKANLEN-CONH_2_, Pepceuticals) and equilibrated for 10 min at room temperature prior to IM-MS analysis. Ion mobility-mass spectrometry data was acquired on a Waters Synapt G2-S*i* instrument operated in ‘resolution’ mode. Proteins were subject to nano-electrospray ionization (nESI) in positive ion mode (at ∼2 kV) with a pulled nanospray tip (World Precision Instruments 1B100-3) prepared as detailed in (48). Ions of interest were mass selected in the quadrupole prior to IMS. The pressure in the TWIMS cell was set at 2.78 mbar (nitrogen), with an IM wave height of 23 V, a wave velocity of 496 m/s and a trap bias of 33.

### Collision-induced unfolding

For collision-induced unfolding (CIU) experiments, the 11+ charge state of Aur A (WT or D274N) in the absence or presence of bound inhibitor was quadrupole-isolated and subjected to collisional activation by applying a CID activation in the ion trap of the TriWave. The activation voltage was increased gradually from 16 to 34 V in two-volt intervals before IMS measurement. CIU was carried out with a travelling wave height of 27 V, velocity of 497 m/s and a trap bias of 35.

### Phosphosite mapping

Purified Aur A was buffer-exchanged into 100 mM ammonium bicarbonate, reduced with 4 mM DTT (30 min, 60 °C), and reduced Cys residues alkylated with 7 mM iodoacetamide (45 min, dark at room temperature), as described previously (49). Proteins were then digested with trypsin (2% (w/w) Promega) for 18 h at 37 °C. RapiGest SF hydrolysis was carried out using 1% TFA (1 h, 37 °C, 400 RPM), prior to LC/MS/MS analysis (49). Phosphopeptide data was processed using Thermo Proteome Discoverer (2.4) and MASCOT (2.6). Raw mass spectrometry data files were converted to mzML format to enable processing with Proteome Discoverer. Data was searched against a human UniProt Aur A database limited to residues 122-403 or the D274N mutation. Processing settings were set as follows: dynamic modifications – Phospho (S/T/Y), maximum missed cleavages – 2, MS1 tolerance – 10 ppm and MS2 mass tolerance – 0.01 Da.

### CCS Calibration and IM-MS Data Analysis

Calibration of the TriWave device was performed using β-lactoglobulin (Sigma L3908), cytochrome c (Sigma C2506) and bovine serum albumin (Sigma A2153) as previously described (50). All data were processed using MassLynx (v. 4.1) and Matlab (2018a) to determine collision cross section (CCS) values. Gaussian fitting was performed in Matlab (Version R2018a), using code minimally adapted from peakfit.m (51) (see supplementary information for detail). Scatter plots of ^TW^CCS _N2>He_ (nm^2^) values versus CCS distribution (CCSD) (nm^2^) were generated using ggplot in RStudio. CIU unfolding plots were generated using CIUSuite 2 (52).

### Western Blotting

Western blotting was carried out using standard procedures. Nitrocellulose membranes were blocked in 5% milk powder (Marvel) in Tris-buffered saline and 0.1% Tween 20 (TBST) (20 mM Tris pH 7.6, 137 mM NaCl, 0.1% Tween-20 (v/v)) for 1 h at room temperature on a shaking rocker. All antibodies were prepared in 5% milk TBST. Anti-phospho Aur A (T288) (Cell Signalling Technologies 2914) was used at 1:5000 dilution and incubated with the membrane for 18 h at 4 °C, as described previously (53). Secondary anti-rabbit antibody (1:5000) was incubated for 1 h at room temperature. X-ray film was exposed to the membrane following application of Immobilon Western Chemiluminescent HRP Substrate (Millipore) developing reagent. The films were developed using an ECOMAX X-ray film processor (Protec).

### Protein kinase activity assays

*In vitro* peptide-based Aur A assays were carried out using a Caliper LapChip EZ Reader platform (Perkin Elmer), which monitors real-time phosphorylation-induced changes in the mobility of a fluorescently labelled Kemptide peptide substrate (5’-FAM-LRRASLG-CO_NH2_) (54). The activity of both WT and D274N Aur A variants (10 ng) was evaluated by incubation with 1 mM ATP and phosphorylation of 2 µM fluorescent peptide substrate in 50 mM HEPES (pH 7.4), 0.015% (v/v) Brij-35, 1 mM DTT and 5 mM MgCl_2_. The activity of Aur A after incubation with TPX2 peptide (5 µM) was determined using a TPX2 concentration range of 0.0004 - 40 µM. To confirm loss of catalytic activity, D274N Aur A was also assayed with 40 µM TPX2 peptide. Data was plotted as % peptide conversion (phosphorylation) over a linear real-time scale, using GraphPad Prism software as described in (55).

### Differential Scanning Fluorimetry (DSF) assays

Thermal shift assays were performed using a StepOnePlus Real-Time PCR machine (Life Technologies) with Sypro-Orange dye (Invitrogen) and thermal ramping (0.3 °C per min between 25 and 94 °C). All proteins were diluted to 5 μM in 50 mM Tris– HCl (pH 7.4) and 100 mM NaCl in the presence of 40 µM inhibitor [or 4% (v/v) DMSO as vehicle control concentration], 1 mM ATP and/or 10 mM MgCl_2_. Data was processed using the Boltzmann equation to generate sigmoidal denaturation curves, and average *T*_m_/Δ*T*_m_ values were calculated using GraphPad Prism software, as previously described (56).

### Molecular modeling

Missing parts of the Aur A sequence were modelled into crystal structures using the PyMod plugin (57) in PyMOL (58). Homology models of the full 122–403 catalytic domain sequence (equivalent to the catalytic domain) were built using MODELLER (59) based on the following Aur A crystal structures: 1MUO, 1OL5, 1OL6, 1OL7, 2WTV (chain A and B), 3E5A, 4C3P, 4CEG, 4J8M, 4JBQ, 5EW9, 5G1X, 5L8K, 5ODT, 6HJK. Where present, phosphorylated residues and the D274N mutation were accounted for, but otherwise the amino acid sequence was the same as UniProt human Aur A accession O14965. All other bound proteins or ligands were removed. The MODELLER loop modelling function in PyMod was then used to build ten improved Aur A models, allowing only the newly added residues of the N- and C-termini (and any new A-loop residues) to change. The model with the lowest ‘objective function’ and without obvious new contacts made with the rest of the protein, was chosen as the starting structure for modelling (30). All-atom simulations were performed with the CHARMM36m force field (60) using NAMD (61). Inputs for NAMD simulations were generated using CHARMM-GUI (62) based on the PYMOD generated models. Phosphorylated residues use the doubly-deprotonated Thr patch (THPB). N- and C-termini were uncapped. The protein was solvated in a rectangular waterbox with a minimum distance of 10 Å between the protein and the box edge (∼20,000 TIP3P water molecules). Cl^−^ ions were added to neutralise the protein. Solvated structures were first subjected to 10,000 conjugate gradient energy-minimization steps. Prior to the collection of trajectory data, a heating protocol that raised the temperature of the system from 0 to 300 K over 60,000 steps and a short pre-equilibration at 300 K for 125,000 steps, were used. The time step of 2 fs was used throughout. Trajectory frames were recorded every 5000 steps (10 ps) and simulations ran for >300 ns with temperature controlled at 300 K and pressure at 1 atm using Langevin dynamics.

Gō-like models and potentials were generated from all-atom initial structures using the MMTSB web service (https://mmtsb.org/webservices/gomodel.html) (63, 64). MD simulations of Gō-like models were carried out using Langevin dynamics and the CHARMM package, version 44/45 (65). The timestep was 10 or 15 fs. Simulations across a range of different temperatures were performed to gauge where the unfolding transition occurs then production simulations were performed below this temperature.

Simulation trajectories were processed and analysed using Wordom (66). The protein component of the system was isolated and aligned, and individual trajectory frames extracted for CCS measurements. IMPACT (67) was used to estimate CCS values for protein structures. The default atomic radii and convergence parameters were used for all-atom simulations. For Gō-like models, atomic radii were estimated to be the average distance between each CA atom (3.8 Å). This provided reasonable comparison with the all-atom simulation results. In all cases the raw IMPACT CCS value based on projection approximation (rather than the recalibrated TJM value) was used, as this provided much better comparison with experimental data for Aur A models and for a bovine serum albumen test model. Clustering of Gō-like model conformers was performed using a 15-Å RMSD cut-off value between clusters. Native contact fractions in Gō-like models were calculated as described by (68) using a low temperature (250 K) simulation to define native contacts distances at 80% occupancy.

## Results and Discussion

### WT but not D274N Aur A is hyperphosphorylated and catalytically active

To evaluate the effects of phosphorylation and small molecule binding on the conformational landscape, dynamics and flexibility of Aur A, we expressed and purified two well-studied Aur A catalytic domain (amino acids 122-403) variants from *E. coli*: a wild-type (WT) active version that extensively auto-phosphorylates during exogenous expression (and exhibits reduced electrophoretic mobility dependent on phosphorylation during SDS-PAGE), and a non-catalytically active variant (D274N), in which the essential DFG motif Asp is replaced with Asn (Supp. Fig. 1A) (20). MS/MS-based phosphorylation site mapping of tryptic peptides from *E. coli* expressed WT Aur A (122-403) revealed at least 6 sites of autophosphorylation (Fig. 1A), including Thr288, which lies in the kinase activation loop and is the classical biomarker for Aur A catalytic activity (23, 49, 69, 70). No auto-phosphorylation sites were observed in D274N Aur A, and this was confirmed by immunoblotting with a phosphospecific antibody against pThr288 (Supp. Fig. 1B). Evaluation of enzymatic activity of WT and D274N Aur A, confirmed that WT, but not the D274N variant of Aur A, exhibited robust catalytic activity towards the substrate peptide in the presence of ATP and Mg^2+^ (Fig. 1B).

**Figure 1.**
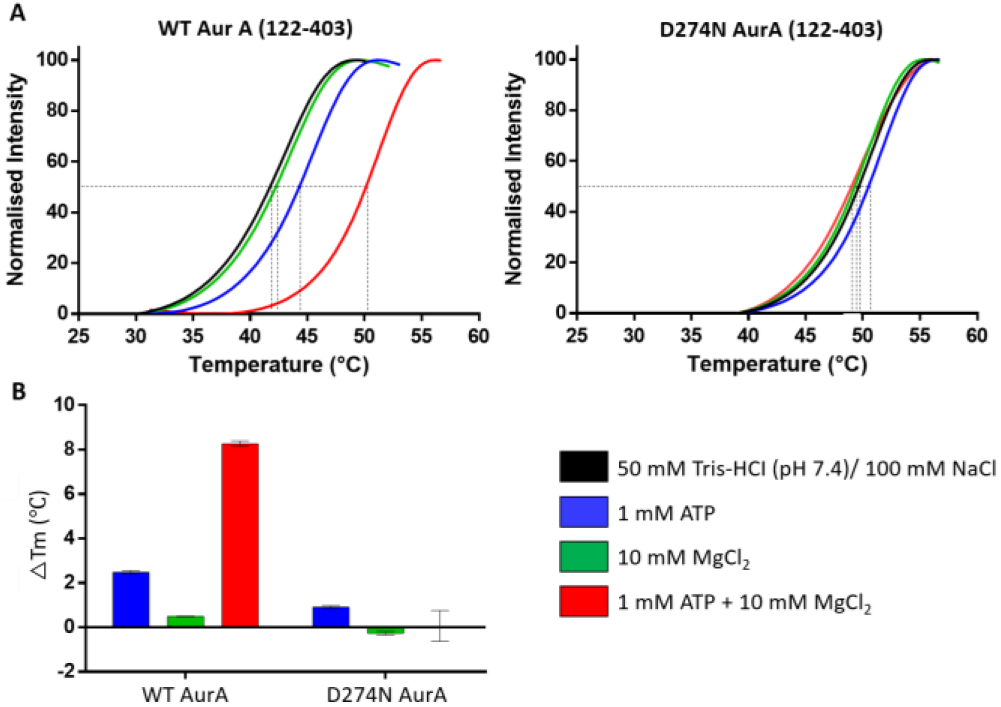
Wild-type (WT) hyperphosphorylated Aur A (122-403) is less thermodynamically stable than a catalytically inactive non-phosphorylated D274N Aur A (122-403) variant. A) DSF thermal stability assay with 5 µM Aur A (black), in the presence of 1 mM ATP (blue), 10 mM MgCl_2_ (green), or 1 mM ATP + 10 mM MgCl_2_ (red). B) Difference in melting temperature (ΔTm) compared with buffer control is presented for both WT and D274N Aur A (122-403).

The thermal unfolding profile of WT Aur A, reported as the *T*_m_ value measured by differential scanning fluorimetry (DSF), increased markedly in the presence of Mg^2+^/ATP (+8.3 °C), which is indicative of tight ATP binding, as previously reported (71). In contrast, there was negligible change (+0.1 °C) in the calculated *T*_m_ of D274N Aur A under the same conditions (Fig. 1C, D), consistent with the inability of this protein to co-ordinate Mg^2+^/ATP. Stabilisation was greatly reduced for the WT protein in the presence of ATP alone (Δ*T*_m_ = +2.5 °C), which is supportive of previous studies showing that Mg^2+^ is required for high-affinity binding of ATP to Aur A (71). WT Aur A demonstrated a lower melting temperature in comparison to D274N Aur A, suggesting that the inactive D274N Aur A protein is also more stable than the active WT form (Fig. 1C, D).

### Active hyperphosphorylated Aur A is less stable, and less conformationally dynamic than the inactive enzyme

To assess the effects of phosphorylation on the structure and conformational flexibility of Aur A, we analysed the hyperphosphorylated WT and the non-phosphorylated D274N proteins by native IM-MS, using travelling-wave ion mobility spectrometry (TWIMS) to determine the rotationally averaged collision cross-section (^TW^CCS _N2→He_) following drift time calibration.

The charge state distribution of both WT and D274N Aur A following native MS was relatively compact (Fig. 2A, B), with 11+ and 12+ charge states of WT Aur A being observed predominantly. The 11+ ion was preferentially observed for D274N Aur A (Fig. 2B). IM-MS analysis of the major 11+ charge state yielded a broad ^TW^CCS _N2→He_ distribution for both protein species (Fig. 2C, D), with the weighted average CCS value for non-phosphorylated Aur A being marginally smaller (22.3 nm^2^) than that for the active phosphorylated Aur A (23.9 nm^2^) kinase. However, the half-height width of the CCS distribution (CCSD) of inactive D274N Aur A was notably broader than that observed for the active enzyme, indicating greater conformational flexibility of the non-phosphorylated protein (Fig. 2E).

**Figure 2.**
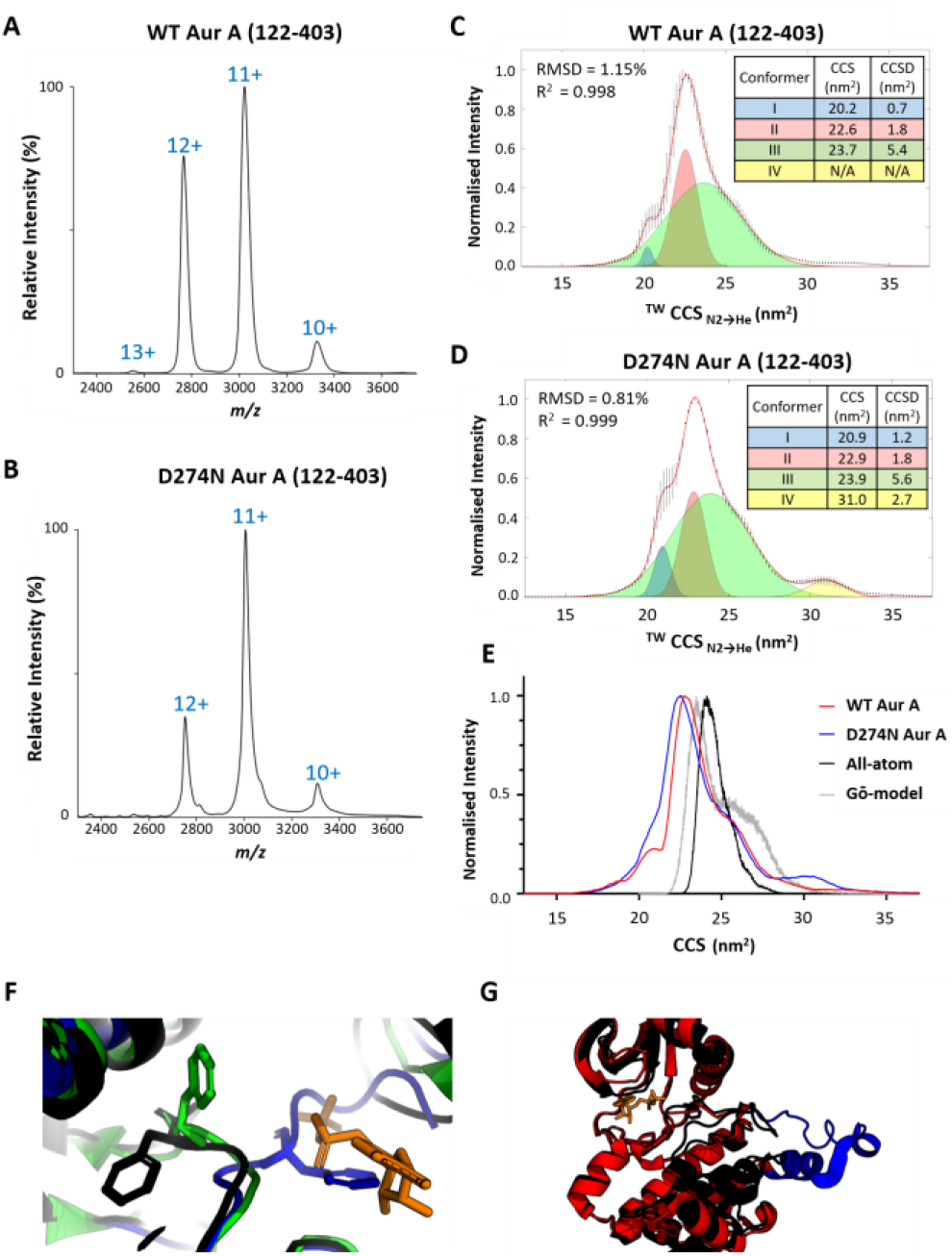
Active hyperphosphorylated Aur A (122-403) is more conformationally compact than inactive non-phosphorylated Aur A. Native ESI mass spectrum of hyperphosphorylated active WT (**A**) or non-phosphorylated inactive D274N (**B**) Aur A (122-403). (**C-E**) ^TW^CCS_N2→He_ for the [M+11H] ^11+^ species of WT (**C**) or D274N (**D**) Aur A (122-403). The red line is the average of three independent replicates. Black error bars representing S.D.. Gaussian fitting was performed in Matlab (Version R2018a), with RMSD and R^2^ values listed. (**E**) Overlaid ^TW^CCS_N2→He_ for WT (red), D274N (blue) Aur A and an overall distribution from all-atom simulations (black). (**F**) Zoomed-in view showing the position of the Phe side-chain in select example crystal structures: DFG-in (black, 1OL7), DFG-up (green, 5L8K), DFG-out (blue, 6HJK). The ATP binding site is marked by an ADP molecule (orange) from 1OL7; this highlights the clash with the DFG-out Phe. **(G)**Overlay of crystal structures from 1OL7 (black) and 4C3P (red). Each 4C3P Aur A monomer exhibits a displaced A-loop and αEF helix (coloured blue) compared to other Aur A crystal structures.

Gaussian fitting of these CCS data revealed three overlapping conformers, with an additional fourth, larger conformational state of relatively low abundance that was fitted for D274N Aur A. For ease of comparison, these data are also presented as weighted distributions for each conformer (Supp. Fig. 2). The CCS and the CCSD values of the two predominant conformers (conformers II and III) for both proteins were within the 2% variance generally observed with these types of native IM-MS experiments (conformer II: CCS = 22.6 nm^2^, CCDS = 1.8 nm^2^; conformer III: CCS = 23.7 nm^2^, CCDS = 5.4 nm^2^), suggesting that these conformational states are likely to be analogous between active phosphorylated and inactive non-phosphorylated Aur A proteins. However, there was a clear difference in the relative abundance of conformers II and III for these two proteins, with the proportion of conformer II being much lower for the non-phosphorylated D274N protein. Conformer I was slightly bigger for D274N Aur A than for phosphorylated WT protein (CCS = 20.2 nm^2^ and 20.9 nm^2^ for the WT and D274N proteins, respectively), and was also a much more abundant component of the conformational space adopted by inactive Aur A. Conversely, the relative abundance of conformer II with respect to conformers I and III was lower in the inactive protein. Instead, we observed an additional configuration, conformer IV at 31.0 nm^2^, exclusively for non-phosphorylated D274N Aur A. These initial observations suggest that phosphorylation of Aur A serves to partially constrain the conformational landscape that this protein can adopt, which is likely linked to successful substrate binding and phosphotransfer.

### Modelling suggests that the major experimental conformers relate to open/closed states rather than different DFG motif conformations or activity

In support of our experimental data, we used IMPACT to estimate CCS values from several all-atom models of Aur A (122-403), built to sympathetically add in missing parts of the protein chain to crystal structures of Aur A found in the protein databank (PDB). These structures include PDB codes 1OL7, 5L8K, and 6HJK, as examples of different configurations of the activation loop (A-loop) with the DFG motif positioned as DFG-in, DFG-up, and DFG-out, respectively (Fig. 2F, Supp. Table 1). One further modelled structure of note was PDB code 4C3P, where dephosphorylated Aur A was co-crystallised as a dimer in the presence of the activating TPX2 peptide, and in which the A-loop and the αEF helix adopt an ‘open’ configuration, extending out from the rest of the kinase domain, and the DFG motif is positioned as DFG-in. Despite the difference in DFG/A-loop position, estimates of CCS values for these static structures—barring those for 4C3P—gave similar values, averaging 22.7, 22.9 and 23.3 nm^2^ for the DFG-in, DFG-up, and DFG-out groups of structures, respectively (Supp. Table 1). These values are an excellent match for those determined experimentally for conformer II or conformer III. The two 4C3P-derived models, with their ‘open’ A-loop structure, gave higher CCS values averaging 24.9 nm^2^, in line with conformer III.

To generate a dynamic picture of protein behaviour, molecular dynamics (MD) simulations were initiated from each of these structural models. Each simulation provides an ensemble of structures and yields a wider distribution of CCS values (Supp. Fig. 3). When considered together, the CCS distributions resemble those observed experimentally by IM-MS for conformers II and III, albeit with the two peak positions shifted up to ∼24 nm^2^ and ∼26 nm^2^ (Fig. 2E, Supp. Fig. 3). This difference of 5–10% compared to the experimentally determined values is similar to the differences previously observed between experimental and IMPACT computed CCS values for other proteins (72). The distributions of exemplar DFG-in (1OL7), DFG-inter (5L8K) and DFG-out (6HJK) structures are virtually indistinguishable, whilst the higher CCS values are almost exclusively from the 4C3P (DFG-in, A-loop open) simulations (Supp. Fig. 3B; Supp. Table 1). An analysis of the DFG motif conformation for every frame of the simulation trajectories shows that, in large part, the initial DFG-motif position is maintained within each simulation (Supp. Fig. 4). Further interrogation of simulated structures (Supp. Fig. 5) suggests that experimental conformers II and III do not relate to different DFG-motif conformations. Instead, conformer II represents ‘closed’ kinase configurations, where the A-loop is inward facing, while conformer III represents the ‘open’ configurations, one example being the dislocation of the A-loop and αEF helix as observed in 4C3P. The broad CCSD of conformer III (Fig. 2C/D) suggests that this is likely made up of several different ‘open’ configurations that cannot be resolved, or possibly interconversion of different ‘open’ and ‘closed’ configurations occurring within the timescale of the IMS experiment.

Enhanced conformation sampling at the expense of chemical detail can be achieved through use of a much-simplified, structure-based ‘Gō-model’, where each residue is considered as a single ‘bead’ and the only stabilising interactions are those from contacts made in the initial structure (68). Individual Gō-model simulations based on the 1OL7 crystal structure gave rise to reproducible CCS distributions that again match well with the experimental profile for conformers II and III (Fig. 2E, Supp. Fig 6). Further analyses of the Gō-model simulated structures again suggest that conformer III could be composed of configurations with a mobile and extended A-loop, but also suggest a significant contribution from configurations with a dynamically, unfolding N-terminus.

Interestingly, none of the MD simulations reveal conformations equivalent to either conformer I or conformer IV as observed by IM-MS, suggesting that these extremes in the conformational landscape may be either experimentally-induced structural compaction in the case of conformer I, or a specific configuration of inactive Aur A that is not represented in the protein data bank (conformer IV).

### TPX2 binding alters the conformational landscape of both hyperphosphorylated and non-phosphorylated Aur A

Binding of the minimal TPX2 peptide (1-43) to phosphorylated Aur A (122-403) has previously been shown to stabilise the active conformation of Aur A *in vitro*, interacting with the N-terminal lobe of Aur A (thereby stabilising the position of the C-helix), and secondarily by stabilising the A-loop (73). We thus investigated the effect of a minimal TPX2 peptide that activates Aur A, on the conformational landscape of both the active and inactive forms of Aur A. Binding of the TPX2 peptide (1-43) to WT Aur A, which increased its activity (Fig. 3A), induced marked differences in its conformational landscape (Fig. 3B). Like unbound Aur A, Gaussian fitting of the CCS profile revealed four conformational states, which we termed I*, II*, III*, IV*.

**Figure 3.**
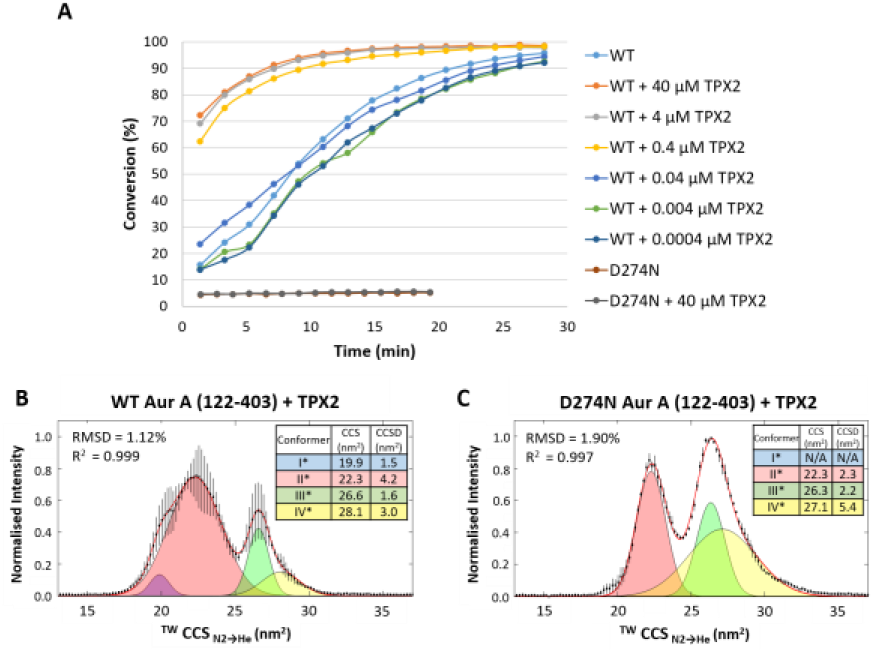
Aur A-activating TPX2 peptide alters the conformational landscape of both hyper- and non-phosphorylated Aur A (122-403). **(A)** *In vitro* peptide-based Aur A kinase assays using 5 µM WT or D274N Aur A in the presence of the minimal activating TPX2 peptide at the indicated concentrations; (**B, C**) ^TW^CCS_N2→He_ of the [M+11H]^11+^ species of WT hyper-phosphorylated active (B) or D274N non-phosphorylated inactive (C) Aur A in the presence of 10-molar excess of the minimal TPX2 peptide. The red line is the average of three independent replicates. Black error bars represent the S.D.. Gaussian fitting was performed in Matlab, with RMSD and R^2^ values listed.

Although the mean weighted CCS values of the two smallest conformational states of WT Aur A are comparable (20.2, 22.6 nm^2^ for I, II respectively, compared with 19.9, 22.3 nm^2^ for I*, II*), the conformational flexibility of these two states notably increased, as defined by approximate doubling of the CCSD values. Further evaluation of conformer II*, and noting the broad CCSD, suggests that this state may be representative of multiple conformations similar to those defined as II and III for unbound WT Aur A (Fig. 2C, Fig. 3B). Given our hypothesis of conformers II and III representing ‘closed’ and ‘open’ configurations of (TPX2 unbound) WT Aur A, these data would suggest that TPX2 binding lowers the barrier for switching between the ‘closed’ and ‘open’ states, changing the conformational equilibrium (and consequently making distinct states harder to distinguish by IM-MS). Furthermore, the fact that there is little difference in the overall CCS between these states in the presence of TPX2 is in agreement with previous studies that reported no global conformation change due to TPX2 (peptide) binding, with TPX2 docking into a (hydrophobic) groove in Aur A (24).

Interestingly, the two larger conformational states, III* and IV*, are distinct from anything observed for either WT or D274N Aur A, although the CCS for III* is similar to that generated following the IMPACT all atom simulation of 4C3P, the TPX2 bound ‘DFG-in’ Aur A where the A-loop is open (Supp. Table 1; Supp. Fig. 3A, B).

Binding of the TPX2 peptide to the inactive non-phosphorylated Aur A yielded two primary CCS distributions, fitting to three Gaussian peaks: II*, III* and IV*; conformer I* was not observed. Supporting our hypothesis that conformer II* for TPX2-WT Aur A represents a dynamic equilibrium between ‘open’ and ‘closed’ conformations, II* for D274N Aur A exhibited a much smaller CCSD (2.2 nm^2^ as opposed to 4.2 nm^2^ for WT Aur A) akin to that seen for unbound protein, as might be expected for inactive protein in a preferentially ‘closed’ configuration.

### Active hyperphosphorylated Aur A is less kinetically stable than inactive non-phosphorylated protein

To better understand relative differences in conformation stability of active hyper-phosphorylated Aur A compared with its inactive counterpart, we performed collision-induced unfolding (CIU) experiments, comparing the CCS of WT versus D274N Aur A at different collision energies (CE) (Fig. 4). The applied CE was sufficient to promote protein unfolding, but not to induce protein fragmentation.

**Figure 4.**
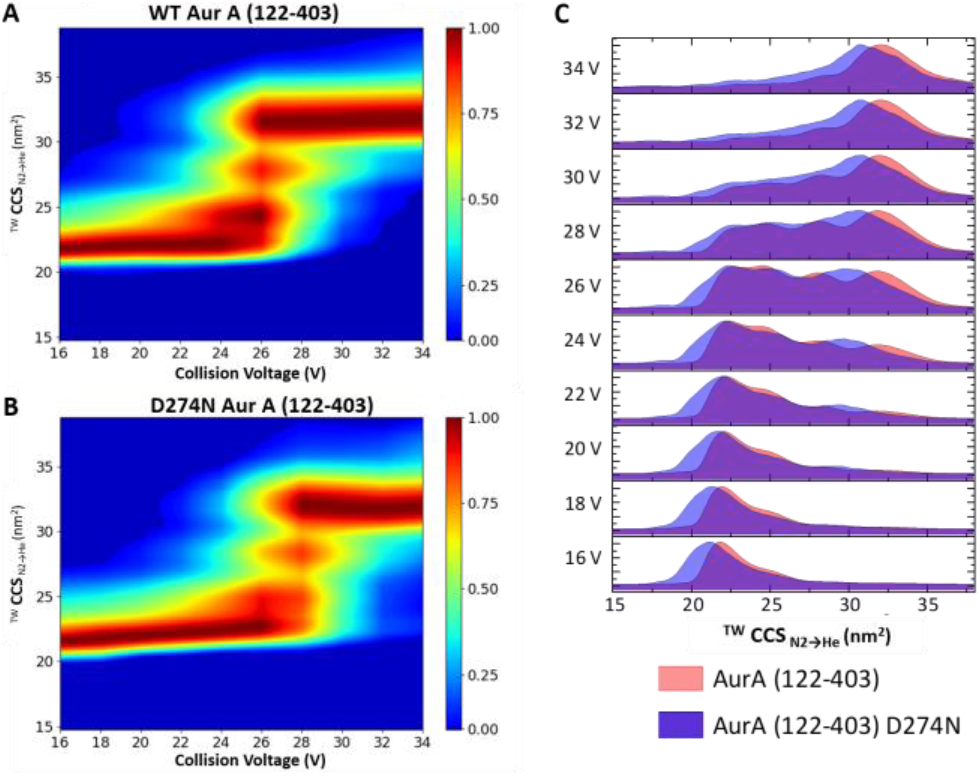
Active Aur A (122-403) is less kinetically stable than inactive Aur A. Collision-induced unfolding profiles for the isolated 11+ charge state of WT (**A**) and D274N (**B**) Aur A (122-403) (or overlaid in (**C**)). Stepped collision energy was applied between 16 and 34 V in two-volt intervals. Data analysis was carried out in MassLynx 4.1, (**A, B**) generating heat-maps using CIUSuite 2 and (**C**) mountain plots using Origin (Version 2016 64Bit). Presented are data from an average of 3 independent experiments.

Comparing CIU profiles in this manner provides information on the relative kinetic stabilities of the two proteins, as the activated ions generated at each stepped CE are trapped in a defined conformational state (54, 74-77). Fig. 4A and B depict the CIU fingerprints for WT and D274N Aur A, respectively, and a direct comparison of the conformational landscapes adopted by these two proteins at each stepped CE value is presented in Fig. 4C. Four main CIU features were observed: the initial conformers (as represented in Fig. 2C, D), two partially unfolded intermediates (ranging from ∼24–28 nm^2^), and final stable ‘unfolded’ states between ∼31-33 nm^2^ for active and inactive Aur A. Similar to the observed differences in conformational space adopted under native conditions, the final ‘unfolded’ inactive non-phosphorylated Aur A had a larger CCSD, indicative of greater conformational flexibility. It is also interesting to note that the CE required to initiate unfolding, and to transition between the partially unfolded intermediates, was lower for active Aur A than was required for D274N (∼24 V versus ∼26 V respectively).

Overall, these data point to the fact that active Aur A is less conformationally dynamic than Aur A in its non-phosphorylated inactive form (albeit with a larger average CCS), and that it is marginally less stable than the inactive protein. This gas-phase kinetic stability data agrees with the liquid-phase thermostability data generated using DSF (Fig. 1C), where the *T*_m_ value (50% unfolding) was 41.7 °C for WT Aur A compared with 49.6 °C for D274N Aur A. Similar findings for solution stability have also been reported elsewhere (78, 79).

### Exposure to small molecule inhibitors alters the conformational distribution of active Aur A

To investigate whether we are able to distinguish modes of small molecule binding to Aur A by IM-MS, we next evaluated the conformational profiles of active and inactive Aur A in the presence of a panel of Aur A inhibitors (Supp. Table 2, Fig. 5; Supp. Fig. 7). Based on a recent analysis, ENMD-2076 should favour a DFG-in mode, whereas MK-8745 is expected to favour a DFG-out mode. MLN8237 and VX-680 are believed to adopt a partial DFG-out position (47). We also investigated the structural effects induced in the presence of the generic type I protein kinase inhibitor staurosporine, which is almost always bound to kinases in a DFG-in conformation. The CCS/CCSD data for each of the (up to) four conformational states, as well as their relative proportion across the conformational landscape (as determined by Gaussian fitting of the CCS profiles for the inhibitor bound Aur A complexes and the proteins alone) are also depicted as proportional plots (Supp. Fig. 7), making differences in the relative abundance of these conformational states (and their relative flexibility) easier to evaluate.

While distinct from unbound active Aur A, the conformational landscapes observed upon binding of each of the Aur A inhibitors was very similar, with comparable CCS and CCSD values for the (up to) 4 conformers defined by Gaussian fitting of the IMS profile. However, comparison with unbound active Aur A revealed a trend towards increased abundance of conformer II relative to conformer III, for all inhibitor-bound forms, as well as a marked increase in the conformational flexibility of conformer I (Fig. 2, Fig. 5, Supp. Fig. 7). Closer inspection of the CCS and CCSD values of conformers II and III across all conditions (Supp. Fig. 7A) reveals some notable differences in conformer III upon inhibitor binding. Not only do the dynamics of this conformer appear to be slightly constrained in the presence of all five small molecules, with a reduction in CCSD from 5.4 nm^2^ to 3.6-4.4 nm^2^, but there is also a ∼5% increase in the average CCS. These data suggest that the inhibitors may help to rebalance the conformational landscape of hyperphosphorylated Aur A towards a population of more discrete states – increasing the relative abundance of the compact ‘closed’ conformer II while decreasing the number of structural permutations underlying the slightly more ‘open’ configuration that we describe as conformer III.

**Figure 5.**
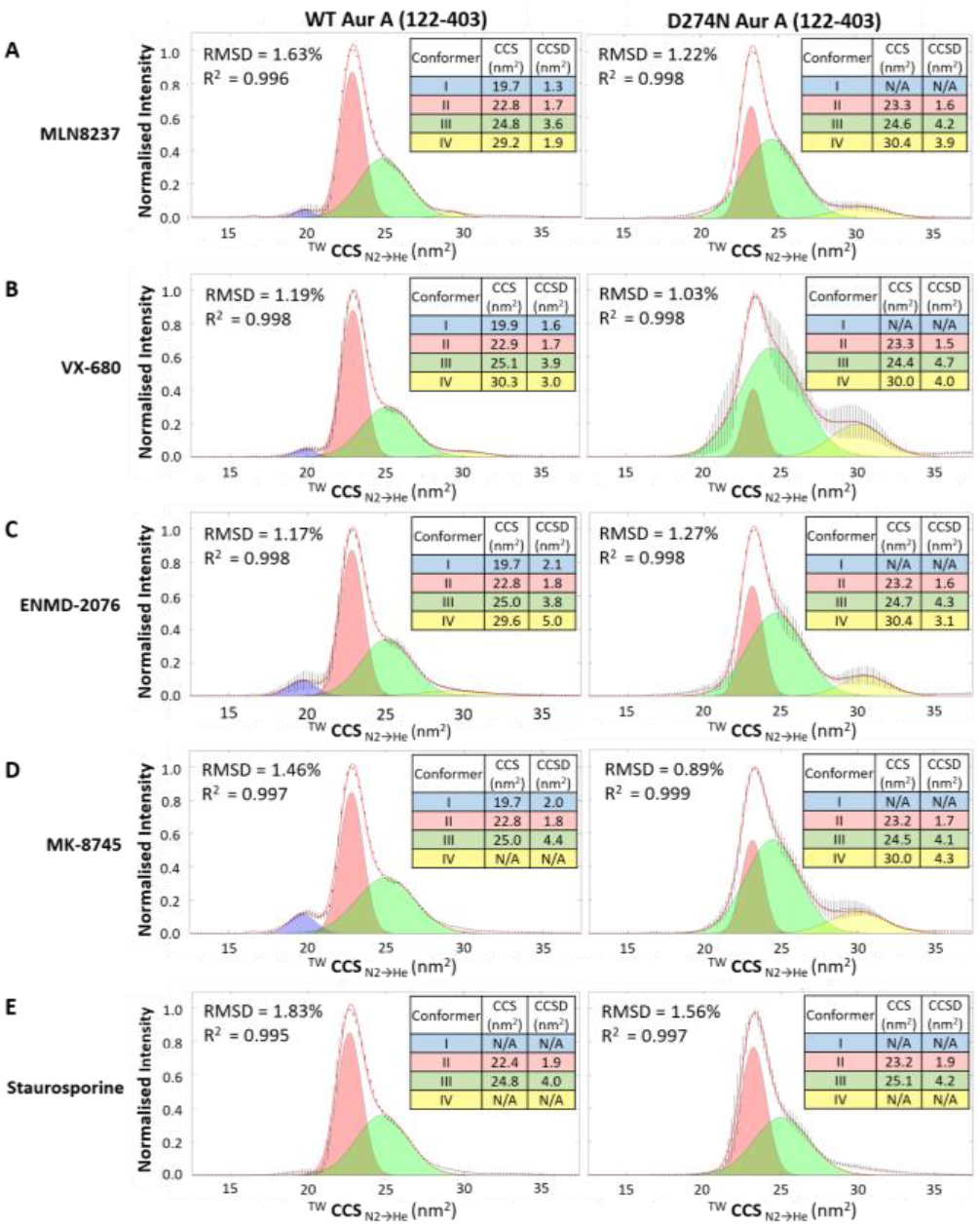
IM-MS of inhibitor bound active and inactive Aurora A (122-403). ^TW^CCS_N2→He_ of the [M+11H] ^11+^ species of WT hyper-phosphorylated active (left) or D274N non-phosphorylated inactive (right) Aur A (122-403) in the presence of 10-molar excess of **(A)** MLN8237, **(B)** VX-680, **(C**) ENMD-2076, **(D)** MK-8745, or **(E)** staurosporine. The red line is the average of three independent replicates. Black error bars represent the S.D.. Gaussian fitting was performed in Matlab, with RMSD and R^2^ values listed.

Comparison of the conformational states adopted by active Aur A bound to either of the two partial ‘DFG-out’ inhibitors, MLN8237 and VX-680 (Fig. 5A, B), as well as the DFG-in ENMD-2076 (Fig. 5C) and DFG-out MK-8745 (Fig. 5D) inhibitors reveal minimal differences. A small increase (3-6%) in the relative abundance of conformer IV at ∼30 nm^2^ was observed for MLN8237, VX-680 and ENMD-2076 (Supp. Fig. 7), whereas evidence for this larger conformational state was absent for the MK-8745-bound protein.

A much greater change was observed in the conformational topology of active Aur A upon binding of the classical non-specific type I (DFG-in) inhibitor staurosporine, with the conformational landscape being constrained to conformers II and III (Fig. 5E, Supp. Fig. 7). Although absent in the presence of staurosporine, the smallest, and generally least-abundant conformational state (conformer I) is present for all other inhibitor-bound complexes of WT Aur A, with a CCS of 19.7-19.9 nm^2^, similar to that observed for the unbound WT enzyme (Fig. 2C) at 20.2 nm^2^.

Interestingly, the conformational landscapes adopted by D274N Aur A in the presence of inhibitors are notably different from each other, and when compared with active Aur A bound to the same small molecule. Of note, the relative abundance of conformer IV increased in all cases, with the exception of staurosporine, where conformer IV is again absent. Conformer I, which accounts for ∼7% of the conformational profile of unbound D274N Aur A was no longer observed, and the ratio of conformers II and III is variable (Fig. 5, Supp. Fig. 7). With regard to staurosporine specifically, and in contrast to the other inhibitors, no difference was observed in the conformational landscape adopted by either WT or D274N Aur A.

### Active Aur A is stabilised to varying extents in the presence of different small molecule inhibitors

The lack of marked differences in the conformational landscapes of active Aur A when bound to the different types of specific Aur A inhibitors prompted us to explore the kinetic stability of these complexes by CIU (Fig. 6; Supp. Figs. 8, 9).

**Figure 6.**
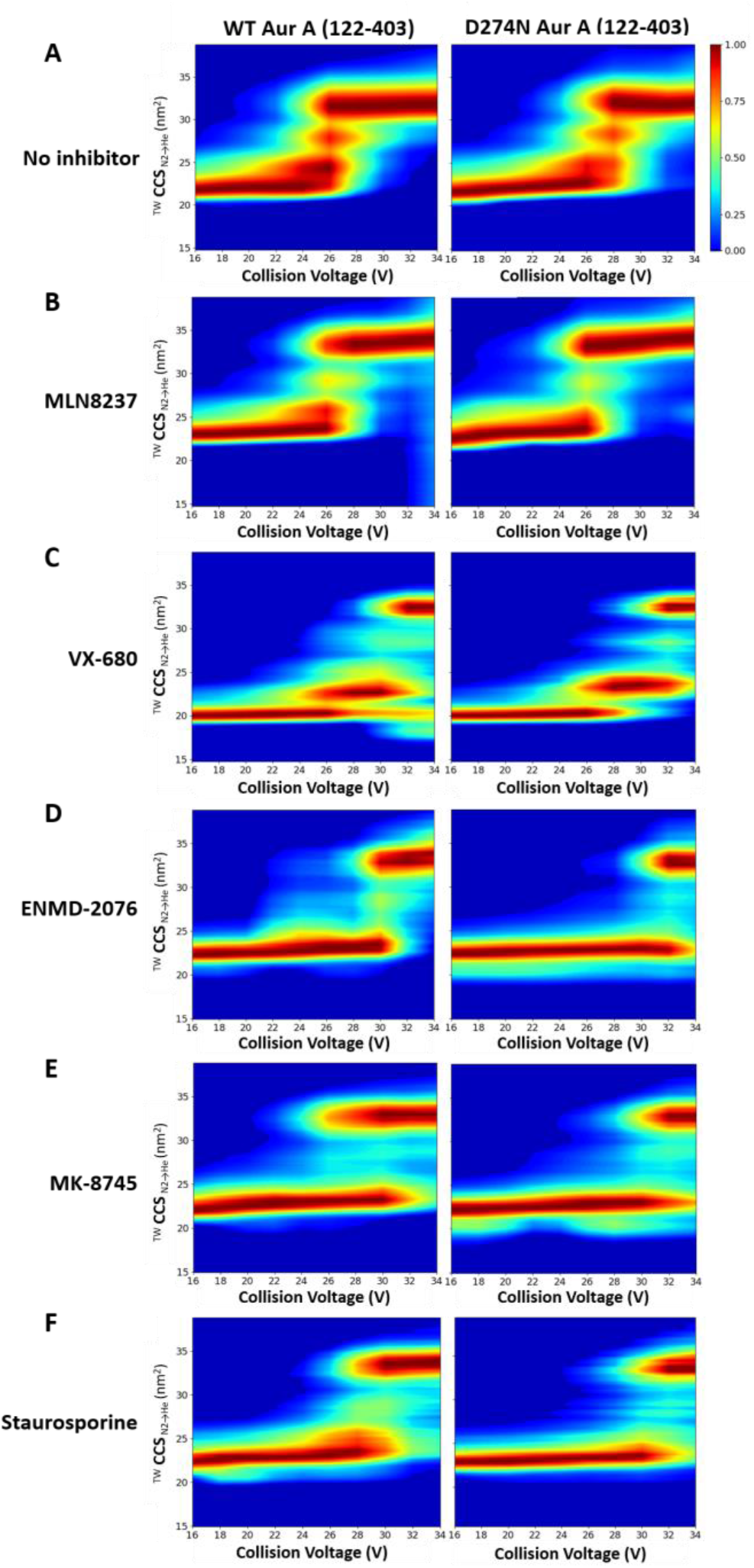
Collision-induced unfolding profiles of inhibitor bound Aur A. The isolated 11+ charge state of **(A)** WT (left) and D274N (right) Aur A (122-403) in the presence of 10-molar excess of **(B)** MLN8237, **(C)** VX-680, **(D)** ENMD-2076, **(E)** MK-8745, or **(F)** staurosporine were subject to CIU using a stepped collision energy between 16 and 34 V (two-volt intervals). Data analysis was carried out in MassLynx 4.1, (generating heat-maps using CIUSuite 2). Presented are data from a single experiment, representative of the data from independent triplicate analyses.

As can be seen from the CIU profiles, the different inhibitors had pronounced effects on the relative kinetic stability of both active and inactive Aur A. While the final stable ‘unfolded’ structures for all the inhibitor-bound forms of WT Aur A (recorded at 34 V) approached a CCS value of ∼ 33–35 nm^2^, the energy required to initiate unfolding, and the conformational states adopted during unfolding were markedly different (Fig. 6, Supp Figs. 8, 9). Of all the inhibitors evaluated, the unfolding profile of MLN8237-bound Aur A (active and inactive) was most similar to that of the unbound protein (Fig. 6, Supp. Fig. 8). MLN8237 had little apparent effect on the kinetic stability of Aur A, as determined by the comparable CE required to induce unfolding. Notably, the CCS of the final unfolded conformation of MLN8237-bound Aur A was larger than for the unbound form, and the relative abundance of the partially unfolded transition states was lower (Fig. 6A, B; Supp. Fig 8), suggesting that the partially unfolded intermediate states were marginally less stable in the presence of MLN8237.

Binding of the other partial DFG-out inhibitor, VX-680, induced a marked stabilisation of the active enzyme, requiring higher CE to initiate unfolding (Fig. 6C). Different (more compact) transition and final states (exhibiting reduced CCSD) were also observed for VX-680–Aur A compared with unbound protein, including a particularly stable partially unfolded intermediate at ∼22.7 nm^2^. There was also some evidence of inhibitor-induced compaction during CIU of WT Aur A, with species of CCS value <20 nm^2^ being observed (Fig. 6; Supp. Fig. 8).

Both ENMD-2076 and MK-8745 transitioned from their native folded state to a stable ‘unfolded’ conformer with limited observable partially unfolded intermediates, albeit with major differences in the kinetic energy required to initiate the process (Fig. 6; Supp. Fig 8). ENMD-2076 induced the greatest stabilisation in WT Aur A, requiring ∼30 V to unfold (Fig. 6D). Although the original conformational states were retained in MK8745-bound Aur A until ∼30 V, transition to the final ‘unfolded’ state was evident by 24 V, with the protein simultaneously adopting two distinct configurations. This difference in unfolding topology for Aur A in the presence of the DFG-out and DFG-in inhibitors was not apparent for the inactive D274N Aur A. Indeed, the CIU profiles for inactive Aur A with either ENMD-2076 or MK-8745 (or staurosporine) were essentially identical.

Comparative thermal stability profiling of unbound versus inhibitor-bound hyperphosphorylated Aur A by DSF (Fig. 7) revealed similar unfolding profiles for WT Aur A in the presence of MLN8237, VX-680 or MK-8745, with an increase in *T*_m_ of >7.5 °C. ENMD-2076 and staurosporine induced slightly greater stabilisation, with a Δ*T*_m_ of >9.2°C. In the case of MLN8237, although we observed thermal stabilisation, there was little difference in the kinetic energy required to initiate unfolding as determined by CIU. However, the CE required to reach the final stable ‘unfolded’ configuration of WT Aur A was higher when it was bound to MLN8237, suggesting that *T*_m_ measurements are likely more representative of the energy required to reach a stable unfolded state. This hypothesis holds true for all inhibitor bound forms of WT Aur A, with the exception of VX-680, but not for inhibitors bound to inactive Aur A. While Δ*T*_m_ values associated with inhibitor-bound D274N Aur A were relatively small (∼<2.5 °C) (Fig. 7), protein unfolding required higher CE for all bound forms. Indeed, with the exception of MK-8745, there was little difference in unfolding profiles, and the CE required to induce unfolding for a given inhibitor, between active hyperphosphorylated, and inactive non-phosphorylated Aur A (Fig. 6, Supp Fig. 8), as exemplified by the comparison of the extracted partially unfolded profiles obtained at a CE of 26 V (Supp. Fig. 9) [78, 80, 81].

**Figure 7.**
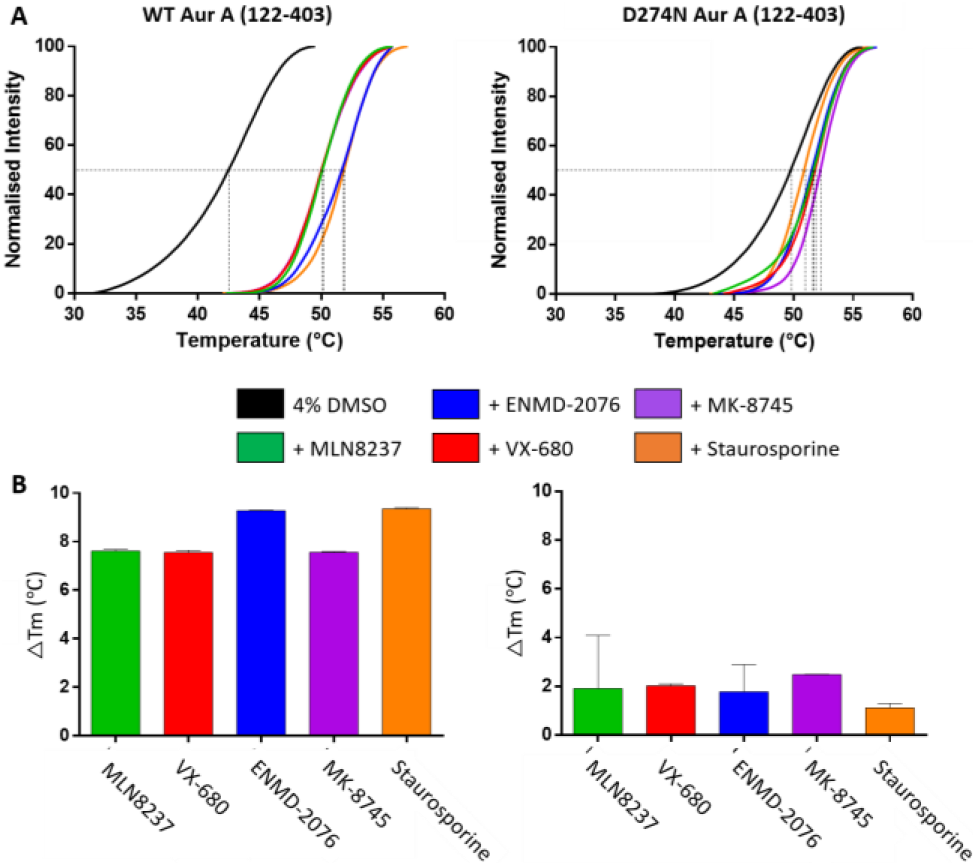
Inhibitor-induced complexation stabilises both catalytically active and inactive Aur A. **(A)** DSF thermal stability assay with 5 µM Aur A + 4% DMSO (black), in the presence of 40 μM of each inhibitor. **(B)** Difference in melting temperature (ΔTm) relative to 4% DMSO control is presented for both WT and D274N Aur A (122-403).

## Discussion and Conclusions

In this study, we exploited IM-MS to explore changes and differences in the conformational landscape dynamics of purified Aur A existing in both active (phosphorylated) and inactive (non-phosphorylated) forms. For the first time, we also examined the effect of an activating TPX2 peptide on Aur A structural dynamics, and evaluated the effects of different classes of small molecule Aur A inhibitors. Gaussian fitting of our IM-MS data reveals up to four conformational states for Aur A, subtle variations in which (such as their relative ratio, CCS and CCSD) are found to be dependent on Aur A activation status, which is previously been shown to correlate with T-loop phosphorylation on Thr288. The active hyperphosphorylated form of Aur A generally exhibits slightly reduced conformational dynamics and reduced stability than the catalytically-inactive protein (Figs. 1 & 2), as determined by both DSF thermal stability and CIU IM-MS experiments. Consistently IM-MS determination of the rotationally averaged collision cross section of active Aur A was further supported by molecular modelling approaches.

Based on evaluation of i) the ratio of conformers II and III across the conditions analysed; ii) the generally broad CCSD of conformer III, suggestive of extensive dynamics, and/or overlapping conformational states that could not be resolved by IMS; iii) the shift to a notably higher CCS and smaller CCSD between conformers III and III* for WT Aur A in the absence and presence of the activating TPX2 peptide; and iv) CCS distributions generated from molecular simulations of Aur A in different conformations being consistent with the experimental profiles for conformers II and III, we propose that conformer II (at ∼23 nm^2^) is representative of ‘closed’ structural states (be they DFG-in/up/out), where the A-loop is inward facing, while conformer III represents one or more ‘open’ kinase configurations where the A-loop extends out.

At the outset of this study, we hypothesised that the activation status of Aur A, and the binding of different classes of small molecule inhibitor would alter the conformational landscape adopted by this protein, as has been established previously (78, 80, 81), e.g. with other protein kinases such as PKA (54, 82), c-Abl (83) and FGFR1 (84), as well as intrinsically disordered proteins such as p53 (13, 85) and Aβ40 (86). While we do indeed see some differences, the effects are subtler than might be anticipated, being broadly consistent with the findings of others (80, 81). Interestingly, all inhibitors increased the ratio of the ‘closed’ conformer II relative to III, with a concomitant increase in the CCS of the third conformational state. However, inhibitor-specific structural effects that have previously been shown to alter the position of the DFG loop using X-ray crystallography, were hard to detect in the gas phase using IMS.

CIU analysis went some way to start to unravel specific inhibitor-induced differences; although all inhibitors stabilised Aur A with respect to unfolding (as confirmed in solution by DSF) and this effect was most marked with the DFG-in inhibitor ENMD-2076, and the partial DFG-up/inter inhibitor VX-680 (Fig. 6, Supp. Fig. 8), as can be seen most clearly when we consider the difference in conformational profiles at mid-unfolding in the snapshot taken at CE of 26 V (Supp. Fig. 9).

Overall, our CIU experiments indicate that all the inhibitors evaluated resulted in kinetic stabilisation of Aur A, given that higher collision energy was required to initiate unfolding, with this effect being least apparent with MLN8237, and highest with ENMD-2076 and VX-680 (Fig. 5, Supp. Fig. 2). More interestingly, by application of CIU we were able to observe differences in the relative kinetic stability of Aur A when bound to the partial DFG-out inhibitors, as opposed to either a DFG-in or a DFG-out inhibitor. Notably, the partially unfolded transition states observed for active Aur A alone or in the presence of either MLN8237 or VX-680 were absent with the other small molecules (Fig. 6) suggesting that these partial DFG-out inhibitors function to ‘lock’ Aur A into specific configurations. The effects of binding of these two partial DFG-out inhibitors to Aur A are thus likely not only a function of the position of the DFG/P-loop, but also reliant on the precise nature of the non-covalent interactions mediated by different chemical classes. We cautiously interpret this finding in the context of the DFG-up conformation, which has been observed with MLN8054 and VX-680 in complex with Aur A (PDB codes 2×81, 2WTV and 4JBQ, respectively). Finally, we anticipate that future studies employing IM-MS and CIU may prove useful in helping to define the mode of action of other small molecule Aur A inhibitors, and in the characterisation of the conformational space adopted by other druggable enzymes, including the >500 members of the human kinome. How this conformational space changes upon binding to a variety of protein and chemical ligands also has potential implications for understanding the different pathways to compound drug-resistance.

## Acknowledgements

L.T. was supported by an MRC DiMeN DTP Ph.D. studentship award (to PAE and CEE). R.B. and M.B. are funded by a CRUK Programme Award (C24461/A23302) and BBSRC Project Award (BB/S00730X/1).

## Author Contributions

L.T., R.B., P.A.E. and C.E.E. designed the project. L.T. performed the majority of the experiments with D.P.B. helping with protein purification, phosphotransferase and DSF assays, J.H. contributing to the MatLab analysis and P.J.B. contributing to mass spectrometry data acquisition. M.B. performed the computational modelling with input from R.B. L.T. and C.E.E. wrote the manuscript with contribution from all authors. All authors have given approval to the final version of the manuscript.

