## Supplementary Material for "Exploring the conformational landscape and stability of Aurora A using ion-mobility mass spectrometry and molecular modelling"

### Supplementary Methods

#### IM-MS Data Analysis – Gaussian peak fitting

Average IM-MS profiles from at least three replicate analyses were used for Gaussian fitting of Aur A conformation states using Matlab (Version R2018a). The code is included below, and was adapted from peakfit.m (O'Hayer T. peakfit.m: MATLAB Central File Exchange; 2021 [Available from: https://www.mathworks.com/matlabcentral/fileexchange/23611-peakfit-m). The maximum number of conformer peaks was set to four, following initial evaluation of up-to six potential conformers, with the final number of conformational states (up to 4) selected based on the minimum error (root-mean-square-deviation (RMSD), $R^2$). Approximate CCS ($nm^2$) and CCSD ($nm^2$) positions were fitted for each conformer using a maximum of 10 iterations with the following starting coordinates: [20,1,23,1.5,25,2.5,29,1.5]; window centre 25.5 units; window width – 19 units; maximum Gaussian width: 1 unit.

*Matlab code:*

```
%data=xlsread('AuroraA_160619.xlsx'); % select the data
signal(1,:)=data([2:199],1)'; % x
signal(2,:)=data([2:199],2)'; % y

RawData=data([2:199],[3:5])';
%SD_RD=std(RawData,0,1); Standard deviation
[R C]=size(RawData);
SD_RD=std(RawData,0,1)/sqrt(R);

plot(signal(1,:),signal(2,:),'k.');
title('Raw Data');  ylim([0 1.1]); xlim([18 32])
 % Input
PeakFit=1; %(1-3,30-33) % see peakfit reference for peak options
https://terpconnect.umd.edu/~toh/spectrum/InteractivePeakFitter.htm#peaksfit
NoP=4; % number of peaks (maximum is 7)
WindC=30; % centre of the window
WindW=25; % width of the window
MinWidth=ones(NoP,1)*1; % minimum width of the gaussians
Iterations=10; % number of attempts to fit the gaussians
Delta=1; % can increase this value if iterations is more than 1
start=[20,1,25,3,29,2]; % set the predicted CCS and CCSD values for each gaussian

figure
plot(signal(1,:),signal(2,:),'k.');
title('Aurora A');  ylim([0 1.5]); xlim([18 34]) % change x and y limit to fit data

[FitResults,FitError]=peakfit_JH(signal,WindC,WindW,NoP,PeakFit,0,Iterations,start);

subplot(2,1,1)
hold on

e=errorbar(signal(1,:),signal(2,:),SD_RD,'k.'); %% for Standard deviation
e.CapSize = 0;
e.MarkerSize=10;
hold off
```

```
title('Aurora A','fontsize',18);
%axes('fontsize',16);
xlabel('TW_C_C_S _N_2_>_H_e_ (nm^2)')
ylabel('Normalised Intensity')
set(gca,'fontsize',16);
set(gcf,'color','w');
x0=10;
y0=10;
width=1000;
height=1000;
set(gcf,'position',[x0,y0,width,height])
txt = ['Error = ', num2str(FitError(1)), ' R^2 = ', num2str(FitError(2))];
t=text(27,1,txt,'fontsize',16);

% In 'FitResults' tab, adjust the CCSD and Area of absent peaks to 0.001 for the colours of peaks to
be consistent throughout whole data set
Fixed=sortrows(FitResults,2);
Fixed(:,1)=[1:4];
% Check Fixed is correct
FaceTran=0.4; % transparency of the shaded area; 0 is invisible and 1 is solid
EdgeTran=0.8; % same as above for the edge of shaded area

% calculates the Gaussian distribution for the parameters given
%FWHM=2.35sigma
s=2.35;
signal(1,:)=data([2:199],1)';
y(1,:) = gaussmf(signal(1,:),[Fixed(1,4)/s Fixed(1,2)]) * Fixed(1,3);
y(2,:) = gaussmf(signal(1,:),[Fixed(2,4)/s Fixed(2,2)]) * Fixed(2,3);
y(3,:) = gaussmf(signal(1,:),[Fixed(3,4)/s Fixed(3,2)]) * Fixed(3,3);
y(4,:) = gaussmf(signal(1,:),[Fixed(4,4)/s Fixed(4,2)]) * Fixed(4,3);

% calculates RMS for the Gaussian plots
Yall=y(1,:)+y(2,:)+y(3,:)+y(4,:);
Diff= signal(2,:)-Yall;
WindStart=WindC-(0.5*WindW);
WindEnd=WindC+(0.5*WindW);
WinXstart=abs(signal(1,:)-WindStart);
[M1 I1]=min(WinXstart);
WinXEnd=abs(signal(1,:)-WindEnd);
[M2 I2]=min(WinXEnd);
Diff=Diff([I1:I2]);
GausErr=rms(Diff)*100;
R = corrcoef(signal(2,:),Yall);
R2 = R(2,1)*R(1,2);

% smooth Gaussians
lengthX = length(signal(1,:));
samplingRateIncrease = 10;
newX = linspace(min(signal(1,:)), max(signal(1,:)), lengthX * samplingRateIncrease);
smoothedY(1,:) = spline(signal(1,:), y(1,:), newX);
```

```
figure
for m=1:4
FaceC=['b','r','g','y','c','m','k'];
smoothedY(m,:) = spline(signal(1,:), y(m,:), newX);
area(newX,smoothedY(m,:),'FaceColor',FaceC(m),'FaceAlpha',FaceTran,'EdgeAlpha',EdgeTran)
hold on
end
%
smoothedYall = smoothedY(1,:)+smoothedY(2,:)+smoothedY(3,:)+smoothedY(4,:);
hold on
plot(newX,smoothedYall,'r','Linewidth',1.5)
xlim([WindStart WindEnd]); ylim([0 1.1]);
%
hold on
e=errorbar(signal(1,:),signal(2,:),SD_RD,'k.'); %% for standard deviation
e.CapSize = 0;
e.MarkerSize=10;

hold off

% change below parameters for figure aesthetics
title(‘Aurora A,'fontsize',18);
xlabel('TW_C_C_S _N_2_>_H_e_ (nm^2)')
ylabel('Normalised Intensity')
set(gca,'fontsize',16);
set(gcf,'color','w');
x0=10;
y0=10;
width=1000; % of the figure window
height=400; % of the figure window
set(gcf,'position',[x0,y0,width,height])
txt = ['Error = ', num2str(GausErr), ' R^2 = ', num2str(R2)];
t=text(27,1,txt,'fontsize',16);
```

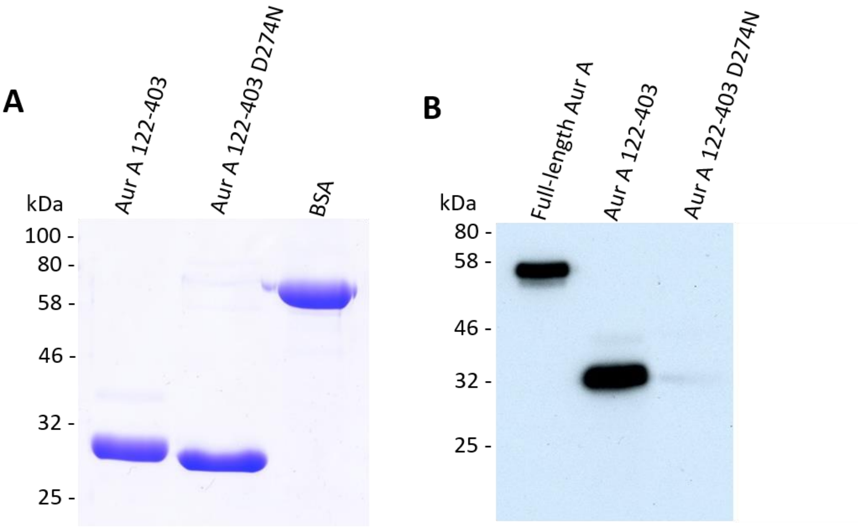

**C**

| Peptide sequence | Modification | Charge | m/z (Da) | DeltaM (ppm) | Ion score | ptmRS |
|---|---|---|---|---|---|---|
| EVEIQsHLR | S186 (Phospho) | 2 | 595.78 | 1.16 | 22 | 100 |
| ELQKLsKFDEQR | S226 (Phospho) | 3 | 534.26 | 1.89 | 32 | 100 |
| DIKPENLLLGsAGELK | S266 (Phospho) | 2 | 888.96 | 0.99 | 61 | 129 |
| IADFGWsVHAPSSR | S278 (Phospho) | 2 | 805.36 | 1.05 | 73 | 100 |
| RttLAGTLDYLPPEmIEGR | T287 (Phospho); T288 (Phospho); M300 (Oxidation) | 2 | 1155.01 | -3.84 | 43 | 99.75; 100 |

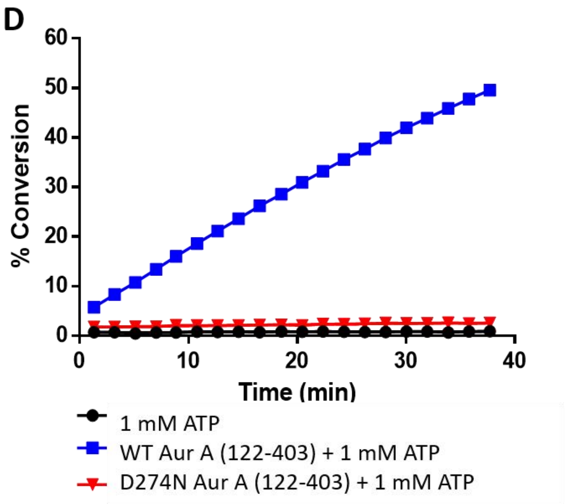

**Supplementary Figure 1. Expression and purification of wild-type (WT) hyperphosphorylated Aur A (122-403) and catalytically inactive D274N Aur A (amino acids 122-403).** (A) SDS-PAGE gel of purified WT and D274N Aur A (122-403) proteins. His-Aur A proteins were purified using a Nickel HisTrap HP column, prior to tag cleavage and gel filtration. Aur A (122-403) has a predicted MW of 32 kDa. BSA was used as a control (B) Immunoblot of full-length Aur A (1-403), WT and D274N Aur A (122-403). Phosphorylation at the activation site in the T-loop (T288) was evaluated using an anti-phospho T288 antibody (CST); 200 ng of protein was loaded per well. (C) LC-MS/MS phosphosite mapping of WT Aur A detailing the peptide sequence identified, the site(s) of modification and the confidence associated with both (Ion score and ptmRS scores respectively) (D) Enzyme activity assay of WT and D274N Aur A (122-403) in the presence of 1 mM ATP (10 ng protein, 2 µM substrate peptide).

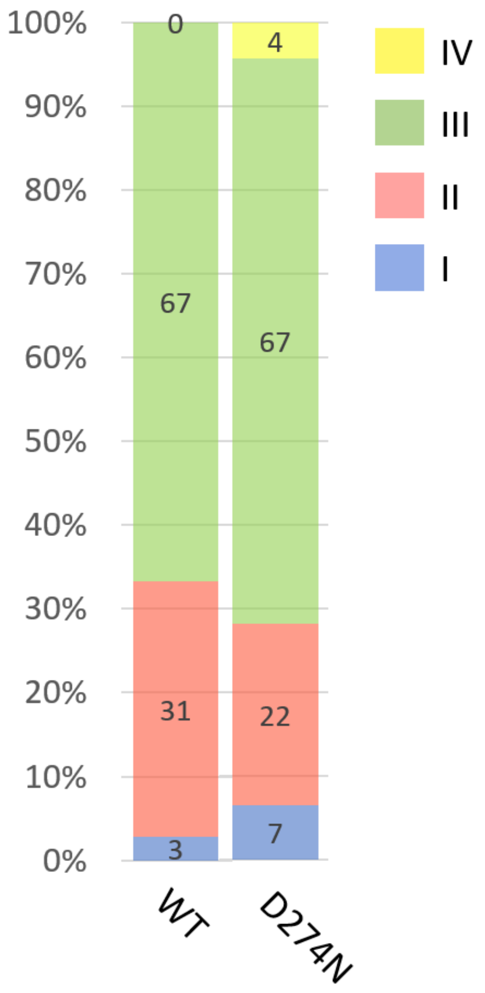

**Supplementary Figure 2. Proportional conformational space adopted by the four different conformers of WT and D274N Aur A (122-403)** (as determined by Gaussian fitting in Fig. 2C and D): I (blue), II (red), III (green), IV (yellow). Average % area presented from three individual experiments.

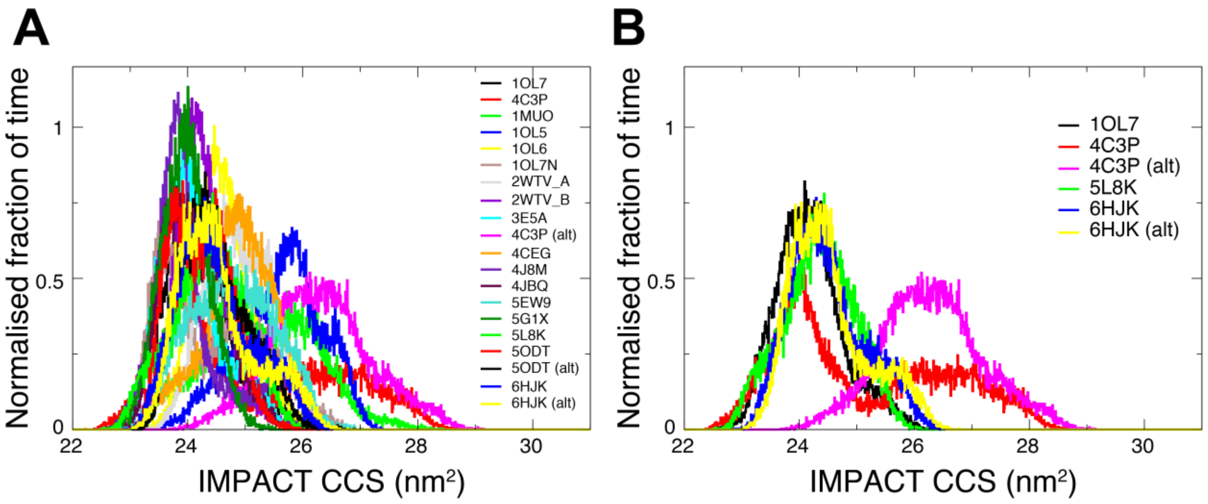

**Supplementary Figure 3. Collision cross section distributions from all-atom simulations of Aur A.** Overlaid normalised distributions for (A) all simulations, and (B) example structures 1OL7 (DFG-in), 5L8K (DFG-up), 6FHK (DFG-out) and 4C3P (DFG-in, A-loop out). The bulk of trajectories contribute to a first major peak at ~24 $nm^2$ and extend between 22.5 and 27 $nm^2$. A handful of models, particularly those of 4C3P, extend the range to higher CCS values contributing to a second peak and the tail of the CCS distribution.

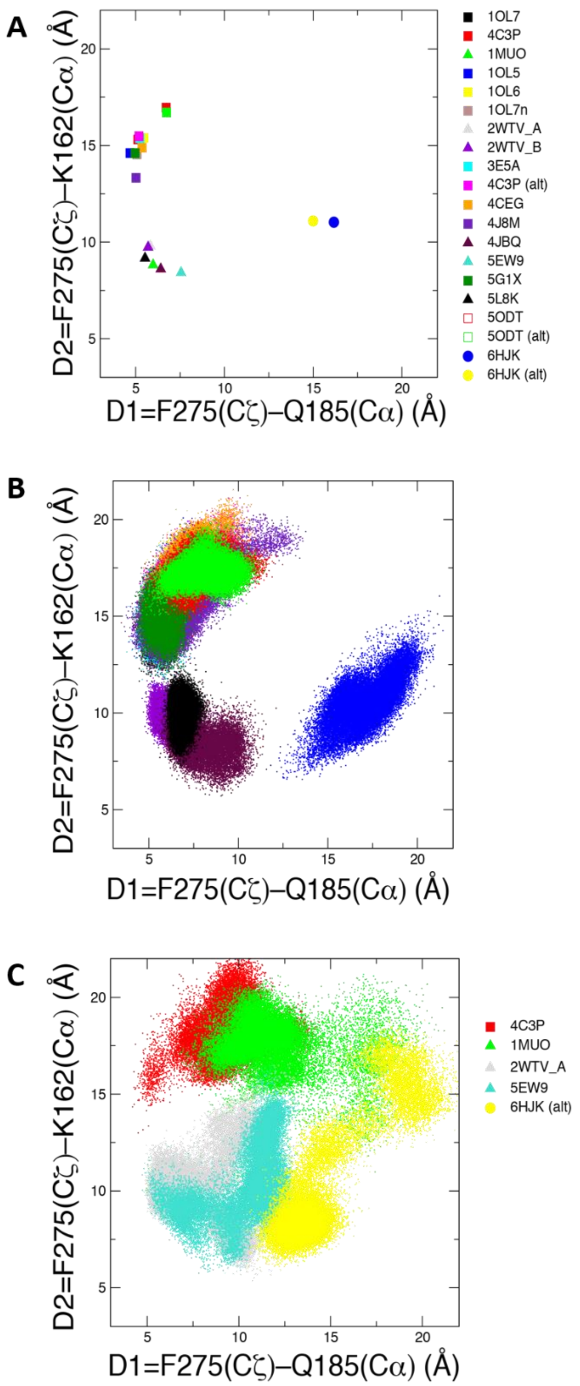

**Supplementary Figure 4. Designation of DFG motif conformations using the D1/D2 criteria introduced by Modi and Dunbrack (PNAS, 2019, 10.1073/pnas.1814279116). (**A) Positions in the D1/D2 plot for each of the starting structures. Three clear groups are observed: the DFG-in group of 12 structures (squares) positioned in the upper left quadrant; the DFG-up/inter group of 6 structures (triangles) in the lower left quadrant; and the two DFG-out 6HJK structures (circles) in the lower right quadrant. (B, C) The range of the D1/D2 plot explored for each simulation. In (B), the majority of simulations remain reasonably close to their initial position and maintain their DFG conformation. In (C), five trajectories are more mobile on the D1/D2 plot and exhibit DFG motif conformations further away from their initial structure, including some conformations in the upper right quadrant of the D1/D2 plot that are not observed in kinase crystal structures.

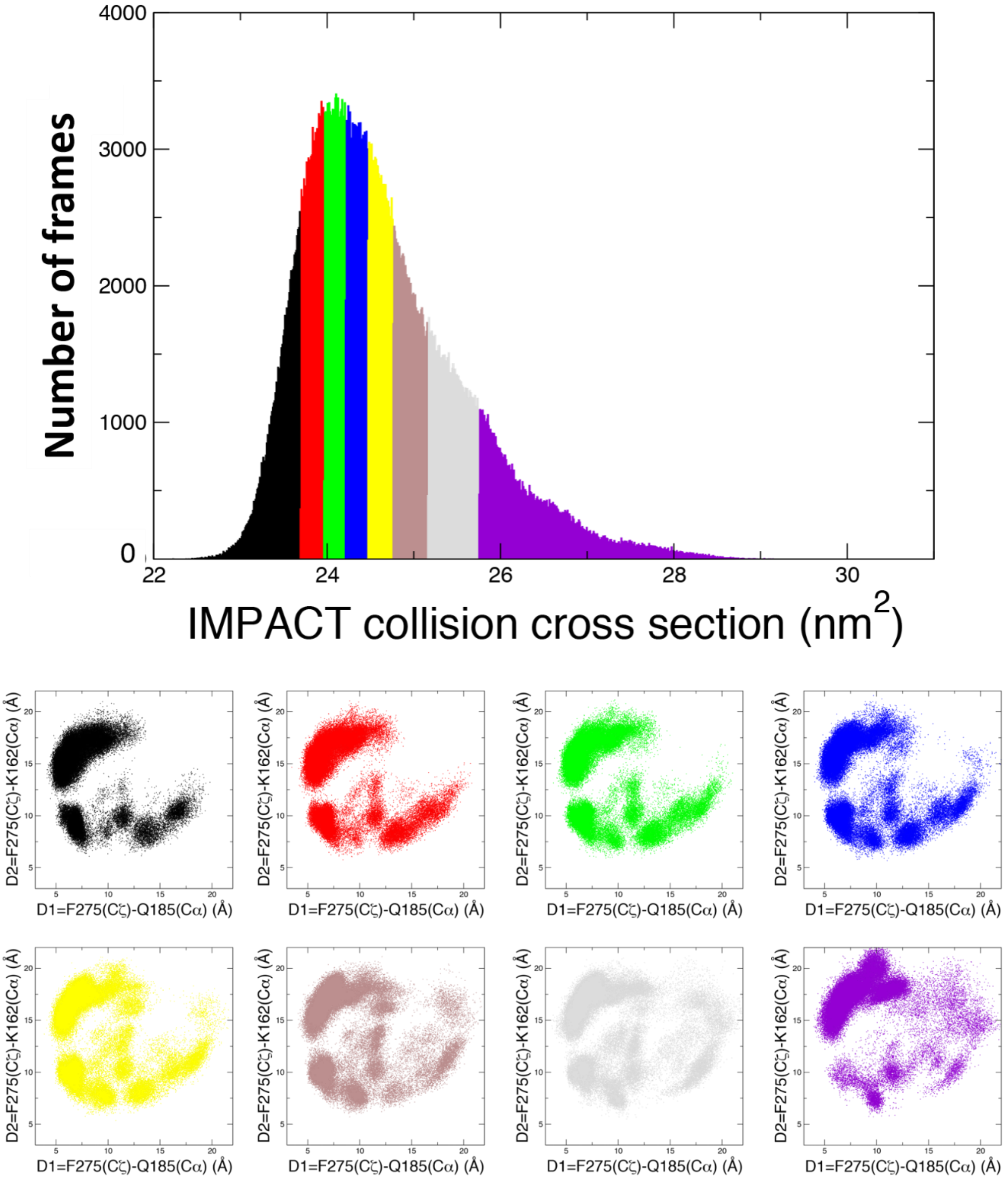

**Supplementary Figure 5. Collision cross section distributions for the combined all-atom simulation trajectories of Aur A.** The distribution has been divided up into eight CCS regions with ~80,000 structures in each region. The contribution of different DFG motif conformations in each region is described using a D1/D2 plot. A high density of DFG-in models is observed in all eight regions on the CCS profile, as shown by the density of structures in the upper left quadrant of the D1/D2 diagram. DFG-in is the most common DFG motif conformer of the initial structures (12 of 20) and includes the A-loop extended 4C3P model. DFG-up/inter and DFG-out conformers are also seen across all eight regions but are less frequently observed at the highest CCS regions. The upper-right quadrant of the D1/D2 plot is more frequently occupied in the highest CCS regions, suggesting that uncategorised DFG motif conformers (not observed in crystal structures) with a more open structure contribute to the high CCS range of the profile.

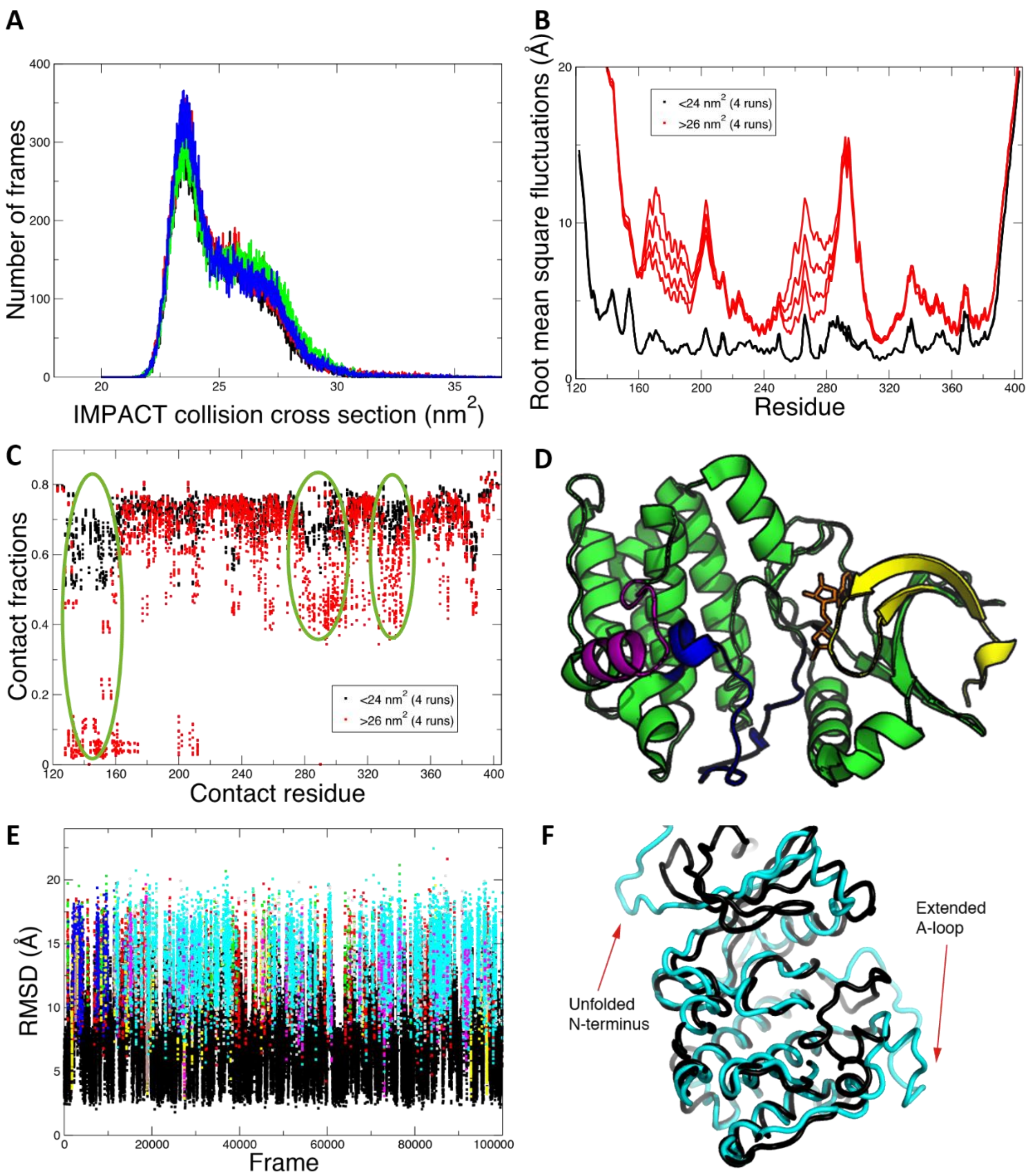

**Supplementary Figure 6. Gō model simulations of Aur A produce CCS profiles similar to those of experimental IM-MS conformers II and III, and suggest that structures with an extended A-loop and unfolded N-terminus may contribute to the higher CCS (conformer III) region.** (A) Overlaid CCS distributions for 4 independent Gō model simulations display the two-component profile akin to those of the combined all-atom simulations and the experimental CCS distributions for conformer II and III. Simulation trajectory frames were separated out into first peak (<24 nm$^2$, mainly ‘conformer II’) and second peak (>26 nm$^2$, mainly ‘conformer III’) groups and then analysed to give: the root mean square fluctuations (RMSF) (B), and the native contact fractions (C). RMSF values are generally higher, marking more variable regions/structures, for the conformer III type (red) ensemble of structures compared to the conformer II-type (black) structures, but especially so in the N-lobe and around the A-loop region. Native contacts are also most likely broken for conformer III-type structures at the N-terminus, and the A-loop and contacting regions, highlighted in (D). An RMSD-based cluster analysis for one of the trajectories, gave representative structures of well-populated clusters with an unfolded N-terminus and extended A-loop region (cyan, E, F).

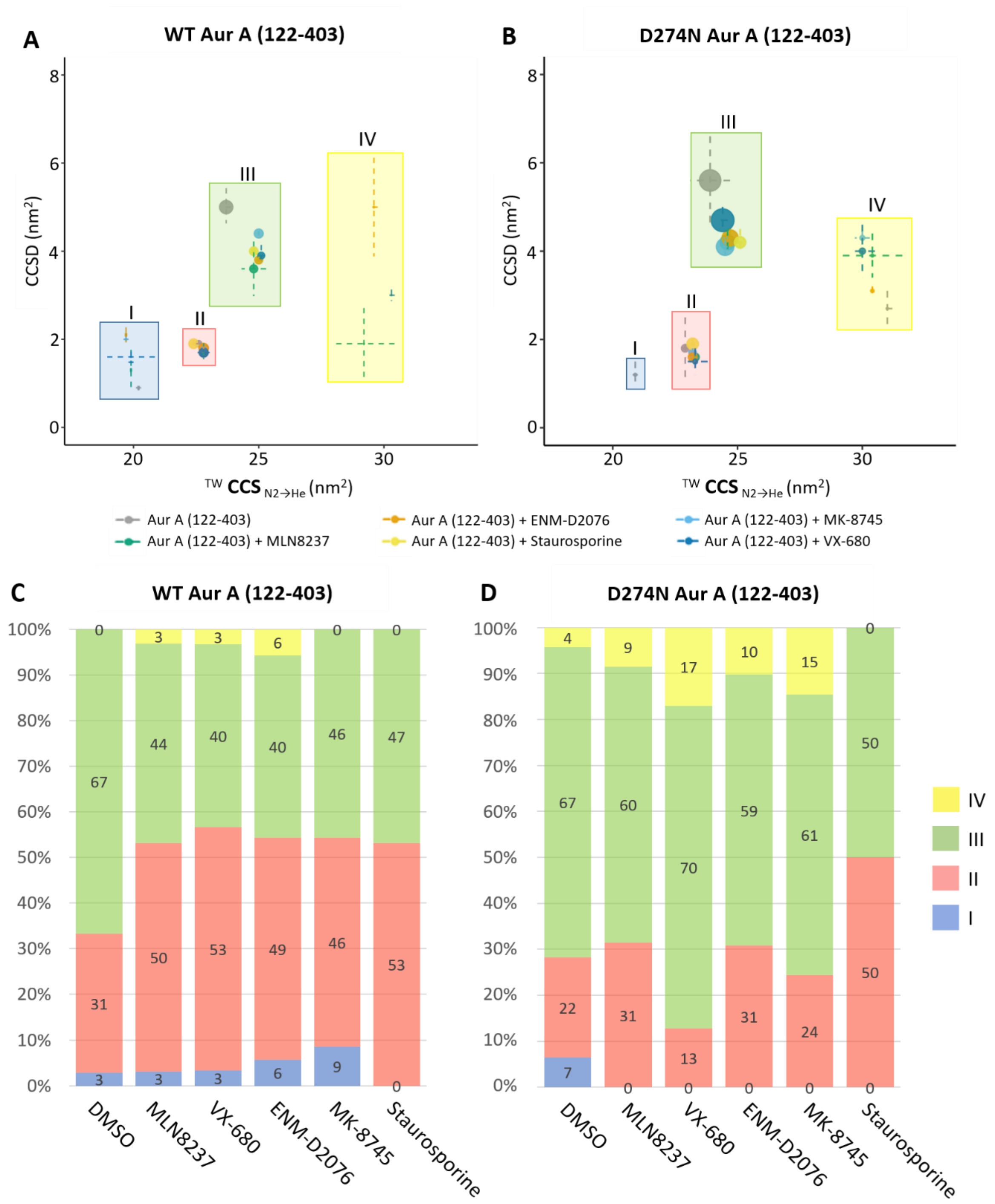

**Supplementary Figure 7. Conformational space adopted by Aur A alone and in the presence of kinase inhibitors reveals inhibitor-mediated stabilisation of the active enzyme.** (A, B) Proportional scatter plots (CCS ($nm^2$) versus CCSD ($nm^2$)) for the different conformational states (as determined by Gaussian fitting in Fig. 5) for WT (A) and D274N (B) Aur A (122-403). Size of dot representative of area. (C, D) % area of four different conformational states (as determined by Gaussian fitting in Fig. 5): I (blue), II (red), III (green), IV (yellow) for WT (C) and D274N (D) Aur A (122-403) alone or in the presence of different inhibitors as indicated. Average % area presented from three individual experiments.

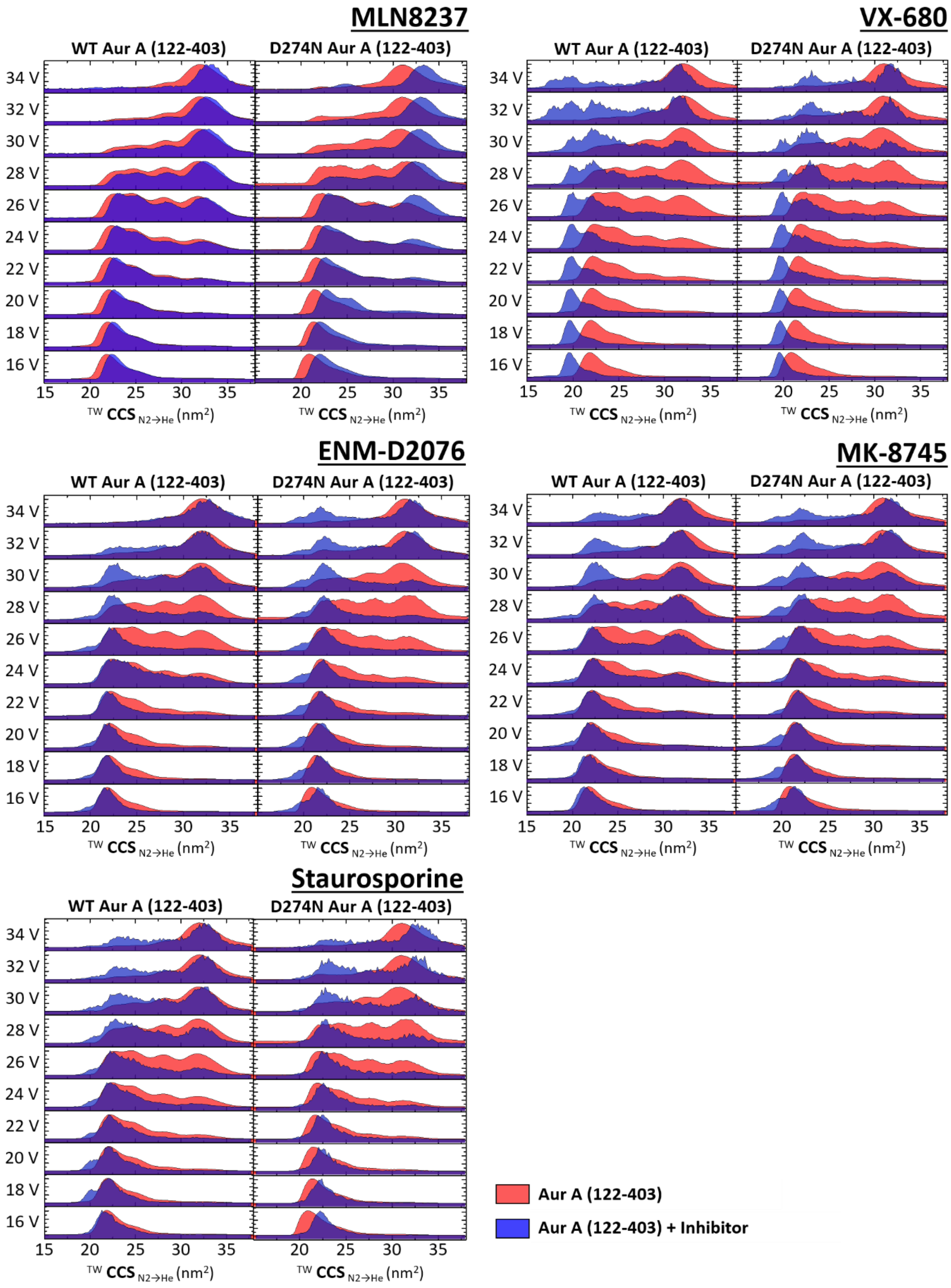

**Supplementary Figure 8. Collision-induced unfolding profiles of Aur A in the absence and presence of inhibitors.** The isolated 11+ charge state of WT (left) and D274N (right) Aur A (122-403) in the presence (blue) or absence (red) of 10-fold molar excess of inhibitor were subject to CIU using a stepped collision energy (CE) between 16 and 34 V (two-volt intervals). Data analysis was carried out in MassLynx 4.1, generating mountain plots using Origin (Version 2016). Presented are data from an average of three replicates.

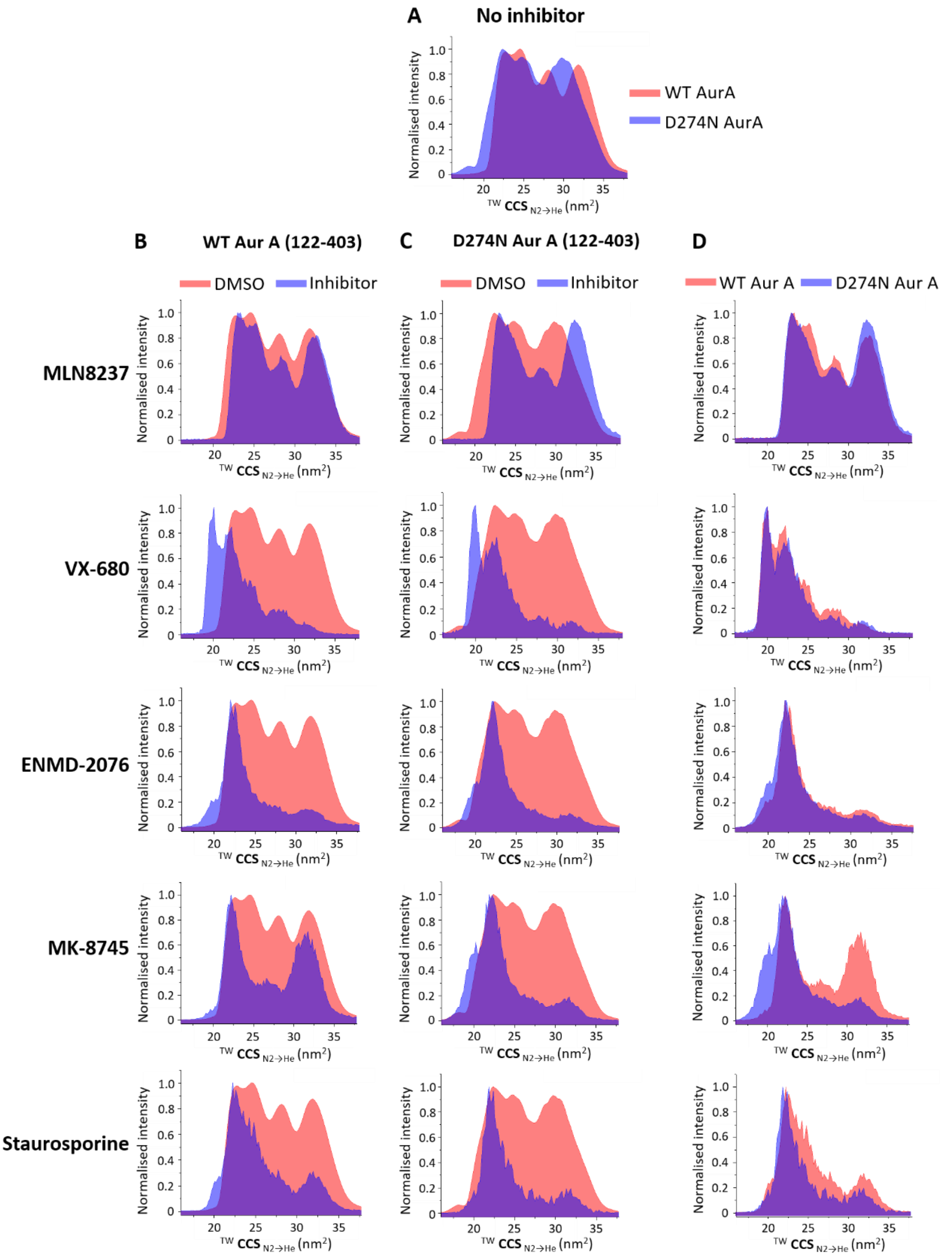

**Supplementary Figure 9. Collision-induced unfolding profiles of Aur A in the absence and presence of inhibitors.** The isolated 11+ charge state at 26 V collision energy of (A) Aur A (122-403) with no inhibitor. (B) WT Aur A (122-403) and (C) D274N Aur A (122-403) in the presence (blue) or absence (red) of 10-molar excess of inhibitor. (D) Aur A (122-403) in the presence of inhibitor for WT (red) and D274N (blue). Data analysis was carried out in MassLynx 4.1 and line plots were generated using Origin (Version 2016). Presented are data from an average of three replicates.

| PDB | [1]CCS ($nm^2$) | Crystal description |
|---|---|---|
| 1OL5 | 22.6–23.0 | phos, TPX2, DFG-in |
| 1OL6 | 22.3–22.8 | dephos, D274N, DFG-in |
| 1OL7 | 22.2–22.7 | phos, DFG-in |
| 1OL7N | 22.1–22.6 | based on 1OL7, dephos, D274N, DFG-in |
| 3E5A | 22.5–22.9 | phos, VX-680, TPX2, DFG-in |
| 4C3P | 25.0–25.5 | dephos, TPX2, dimer, DFG-in, A-loop open |
| 4C3P (alt.) | 24.3–24.8 | dephos, TPX2, dimer, DFG-in, A-loop open |
| 4J8M | 23.1–23.7 | dephos, T287D, CD532, DFG-in |
| 4CEG | 21.1–21.6 | phos, C290A, C393A, DFG-in |
| 5G1X | 21.8–22.2 | phos, N-MYC, DFG-in |
| 5ODT | 23.2–23.7 | dephos, D274N, TACC3, DFG-in |
| 5ODT (alt.) | 23.9–24.3 | dephos, D274N, TACC3, DFG-in |
| 1MUO | 23.2–23.6 | dephos, adenosine, DFG-up |
| 2WTV | 23.4–23.7 | phos, MLN8054, chain A, DFG-up |
| 2WTV | 22.6–23.0 | phos, MLN8054, chain B, DFG-up |
| 4JBQ | 21.9–22.5 | dephos, VX-680, DFG-up |
| 5EW9 | 22.8–23.4 | phos, MK-5108, DFG-up |
| 5L8K | 22.3–22.7 | phos, vNAR, DFG-up |
| 6HJK | 22.9–23.4 | dephos, L210C, ASDO2, DFG-out |
| 6HJK (alt.) | 23.0–23.4 | dephos, L210C, ASDO2, DFG-out |
| 4F5S | 39.3–40.1 | BSA, calibrant |

**Supplementary Table 1. Collision cross section estimates made using IMPACT on a single initial structural model of Aur A (122–403) for each of the listed PDB codes.** DFG-in models are grouped in blue, DFG-up (or inter) models are grouped in yellow, and two alternatively built starting structures for the only available DFG-out Aur A structure (6HJK) are green.

[1] As IMPACT is a stochastic method, CCS estimates were performed 200 times on each structure to give a range of values.

| Inhibitor | Mass (Da) | Structure | Binding configuration |
|---|---|---|---|
| MLN8237<br>(Alisertib) | 518.92 | 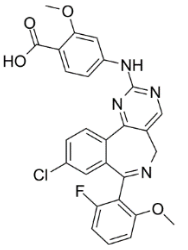 | DFG-out/up<br>PDB 2X81 for very closely-related compound MLN8054 |
| VX-680<br>(Tozasertib) | 464.59 | 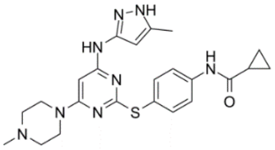 | DFG-in/up (with/without TPX2) |
| ENMD-2076 | 375.47 | 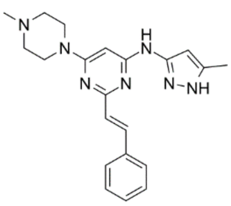 | DFG-in |
| MK-8745 | 431.91 | 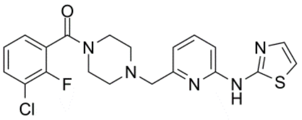 | DFG-out |
| Staurosporine | 466.53 | 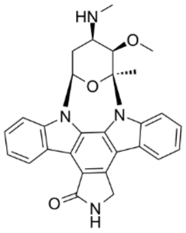 | ND |

**Supplementary Table 2. Aur A inhibitors employed in this study. ND: Not Determined**
